# Genomic history and ecology of local adaptation in sky-island *Arabidopsis thaliana*

**DOI:** 10.64898/2026.05.25.727657

**Authors:** Diana Gamba, Amanda Penn, Abigail Lukasevics, Asnake Haile, Jeffrey Kerby, Mistire Yifru, Tigist Wondimu, Collins E. Bulafu, Jesse R. Lasky

## Abstract

Adaptation to local environments in isolated habitats may produce independent solutions to similar ecological challenges, yielding insight into the predictability of evolution. Alpine plants on tropical sky-islands, in particular, must cope with extreme diurnal fluctuations in temperature and radiation. These populations may undergo parallel local adaptation to high elevation, or alternatively, distinct local adaptations due to intermountain heterogeneity. For instance, abiotic stress on mountain peaks could select for stress tolerance or stress escape/bet-hedging strategies. We tested these hypotheses in experiments with 163 newly sequenced *Arabidopsis thaliana* genotypes from East Africa. Genomic history followed geography: Ugandan genotypes were the most divergent, and Ethiopian genotypes more closely related. Effective population size decreased towards the present on all mountains, in concert with declining climatic suitability since the end of the African Humid Period. Genotypes from the colder and seasonally drier mountains showed strong seed dormancy, rapid flowering, short/curved inflorescences. In contrast, genotypes from the milder and less seasonal mountains showed less seed dormancy, delayed flowering, and long/erect inflorescences. Elevational clines found in multiple mountains included seed dormancy-release with short (vs. long) stratification, and shorter inflorescences, at higher elevations. In a genome-wide association study (GWAS) with inflorescence length, we detected the *TCP5* transcription factor, but these variants only segregated in some mountains and showed an elevational allele frequency cline at only one mountain. Allele reuse based on GWAS was greater between Ethiopian populations (QTL allele-frequency *r* > 0.9 for most traits), and generally lower between Uganda and the rest. Results indicate limited evidence of parallel local adaptation along elevation, suggesting that dwarf inflorescences are selected for in multiple mountain tops, while bet-hedging prevails in cold and seasonally dry sky-islands, and divergent selection along elevation in other traits can evolve in mountains with milder climates. Overall these results suggest a long history of isolation and ecological complexity limit the predictability of local adaptation.

## Introduction

Alpine environments are often isolated and thus have become models for understanding repeated evolution under similar climatic pressures, and key for studying the predictability of evolution^1,2^. Plant communities at high elevation deal with extreme cold, high solar radiation, and wind^3^, often converging phenotypically and specializing on those conditions^3–6^. Similarly, alpine populations of plant species may show reduced plant size and increased leaf thickness compared to lowland populations^7^. Adaptive convergent evolution appears to entail genetic reuse across deep time, from closely to distantly related species, suggesting that genetic adaptation to extreme environments could involve only a few pathways^8,9^. Recent empirical evidence suggests that gene reuse in alpine environments depends on the time scale considered, being more likely in recently diverged lineages (e.g., populations of the same species) than in more anciently diverged ones (e.g., species in the same genus)^10,11^. Mountain environments, however, can be highly heterogeneous, potentially driving population differentiation/incipient speciation via geographic isolation and ecological adaptation^12^. It is less clear if climate heterogeneity among alpine sky-islands promotes diversity of local adaptations within species.

Studies of intraspecific convergent adaptation to alpine environments have generally focused on temperate regions^1,6,7,10,13^, while tropical alpine environments are distinct. They are important hotspots of biodiversity and endemism^4,14–17^ and could harbor diverse local adaptations within species. In these sky-islands plants must adjust to extreme UV radiation and daily fluctuations in temperature (*i.e.*, night-time freezing), as opposed to marked temperature seasonality, displaying great diversity of adaptations across species^4,5,16,17^. In a plant community of the Venezuelan Andes, different plant growth forms segregated along an axis of avoidance-tolerance strategies related to daily fluctuations in temperature and water availability^18–20^. In the flora of East African sky-islands, gigantism (*e.g.*, in *Dendrosenecio, Lobelia*) and dwarfism (*e.g.*, in *Alchemilla*) have sometimes evolved independently in isolated mountains^5,15^. Within species, high-elevation adaptation may thus be conserved, *i.e.*, there is a single tropical alpine strategy. Moreover, phenotypic plasticity and bet-hedging could be critical for persistence in these tropical sky-islands as they are endangered ecosystems with increasingly unpredictable conditions^16,21,22^. Measuring traits associated with life history (*e.g.*, seed dormancy, flowering time, plant size) is thus critical for examining the targets of selection in tropical sky-island adaptation(s).

Phenology, an important component of plant life history, mediates the interplay of development under optimal environmental conditions^23,24^. Natural genetic variation in phenology often underlies local adaptation, shaping life history strategies in different environments^25^. For example, in the model plant *Arabidopsis thaliana* genetic variation in the timing of germination and flowering is adjusted to local seasons across Eurasia^26,27^ and genes controlling flowering time show strong signals of local adaptation^28,29^. Such seasonal timing can be classified in four broad adaptive strategies: Mediterranean rapid cyclers (early flowering and low germination/high primary dormancy in the spring/cold-released dormancy; e.g., lowland Iberia^30^), weedy spring-summer cyclers (early flowering and high germination in the spring/cold-released dormancy and summer; e.g., some urban populations in Europe^31^), facultative winter cyclers (mid to late flowering and low germination in the fall/cold-induced dormancy, or germination in the spring in years or sites with mild winters; e.g. mountainous Iberia^32,33^), and strict winter cyclers (late flowering and intermediate germination in the fall/cold-induced dormancy; e.g. Scandinavia^34^). *Arabidopsis thaliana* is also a model for studying alpine adaptation. For instance, putatively adaptive strategies include late flowering/high water use efficiency in alpine western Mediterranean and early flowering/low water use efficiency in alpine central Asia^35^. While Eurasian alpine adaptation has been well-researched in this model plant^32,36–38^, the native populations in East African tropical sky-islands remain less studied.

The East African populations of *Arabidopsis thaliana* are an ideal system to study adaptation and its genetic bases in tropical alpine ecosystems. These populations of Arabidopsis were recognized for their unique alpine characteristics by Hedberg (1964)^39^, a pioneer of Afroalpine plant ecology and evolution^4,40^. Later researchers erroneously considered these non-native recent introductions^41^, but recent studies show that they are old and can be genetically diverse^35,42–44^. These populations also occupy outlier environments compared to the rest of the species range and might have marginal habitat suitability^45^. Genetic variation across East Africa appears to be highly geographically structured^35,42^, suggesting that sky-island colonization has occurred independently multiple times. Field observations suggest phenotypic convergence in different Afroalpine mountains. However, we know little about natural genetic variation in ecologically important traits or to what extent populations are more sensitive when exposed to different conditions. If sky-island convergent adaptation has evolved in Afroalpine Arabidopsis, we would expect repeated and putatively adaptive trait-elevation clines between mountains, in line with all sky-island populations undergoing selection towards a common optimum (*i.e.*, “global” adaptation^46^). Alternatively, heterogeneous deglaciation histories and present climates^47,48^ could have promoted different alpine strategies where each mountain top is a reservoir of distinct genetic diversity in traits. For instance, in climates with seasonal precipitation, bet-hedging and consequent seed banking could be selected for in mountain peaks with increasingly unpredictable drought, while more conservative stress-tolerance strategies could be selected for in mountain peaks with milder climates.

In this study we integrate multiple lines of evidence to test for adaptation in tropical Afroalpine populations of *Arabidopsis thaliana*. We focused on populations from four isolated mountains: three in the Ethiopian Highlands and one on a large shield volcano in Uganda. We analyzed whole genomes to detect patterns of historical diversity and divergence times among mountain populations. We examined multiple dimensions of seed dormancy and measured genetic variation in flowering time and plant size. We also conducted prolonged cold-exposure experiments to examine genetic sensitivity in the same traits. Finally, we implemented genome-wide association studies (GWAS) and evaluated allele reuse across mountains. We asked:

1. Is adaptive potential different across Afroalpine mountains? Previous evidence suggests independent Afroalpine colonization across East Africa and potentially higher genetic diversity in the Bale Mts.^42^, thus we expect differences in genetic diversity/adaptive potential between mountains.
2. Are populations in different Afroalpine mountains phenotypically similar? We hypothesized that due to steep geographical isolation between peaks, potentially resulting in distinct local selection pressures, populations from different mountains differ phenotypically. Alternatively, similar phenotypes may have evolved due to “global” adaptation (*sensu*^46^) to stressful Afroalpine climates.
3. Is local adaptation along elevation repeated across Afroalpine mountains? We hypothesized that similar conditions at higher elevations have shaped a shared locally adaptive strategy across sky-islands. Alternatively, heterogeneity in genomic and climatic histories could have precluded repeated evolution along elevation at each mountain. For instance, in mountain peaks with extreme environments (*e.g.*, seasonally dry) a bet-hedging strategy (*e.g.*, strong seed dormancy and rapid flowering) might prevail, while in mountain peaks with milder environments (*e.g.*, less seasonal) a stress tolerance strategy (*e.g.*, little seed dormancy and delayed flowering) might be advantageous.
4. Are high-elevation genotypes less sensitive to cold than low-elevation ones? We hypothesized that prolonged cold strengthens seed dormancy, delays flowering time, and reduces overall plant size, but that the highest-elevation genotypes would be less sensitive due to local adaptation. Alternatively, cold effects could be more pronounced (*i.e.*, greater sensitivity) in high-elevation genotypes, suggesting adaptive plasticity.
5. Does genetic similarity shape allele reuse in quantitative trait loci? We hypothesized that closely related mountain populations will show greater allele reuse (*i.e.*, higher allele frequency correlation) in quantitative trait loci detected with GWAS. Alternatively, climatic similarity between mountains may better reflect the extent of allele reuse.

## Results

### Q1. Genomic and climatic history reflects deep heterogeneity between mountains

Genetic variation of Afroalpine *Arabidopsis thaliana* was structured by mountain of origin rather than elevation, indicating that alpine colonization occurred independently in each sky-island, consistent with what we have observed in most (but not all) of Eurasia^35^ (Fig. 1). Using ∼146K unlinked single nucleotide polymorphisms (SNPs; from resequencing with short Illumina reads) from our diversity panel of 163 genotypes, two clustering methods showed that genotypes from different sky-islands belonged to different isolated lineages. Each mountain was mostly composed of a single genetic cluster, except for genotypes from the Bale Mts., which were the best sampled (Fig. 1B, C showing k=6 ancestral genetic clusters). Intermediate genotypes were rare, except for the small number sampled from the Guassa Plateau (n=4 from two sites) which were assigned to a mix of all clusters. Population genetic differentiation was high between mountains (*F_ST_* ranged from 0.191–0.318), with genotypes from the equatorial and southernmost Mt. Elgon being the most differentiated, while the central Bale Mts. being more similar to other mountains (Fig. 1D). Cluster assignment of Bale genotypes largely followed latitude (*i.e.*, isolation-by-distance) (Fig. 1F). Due to small sample size in Guassa, we excluded these genotypes from subsequent analyses.

**Fig. 1.**
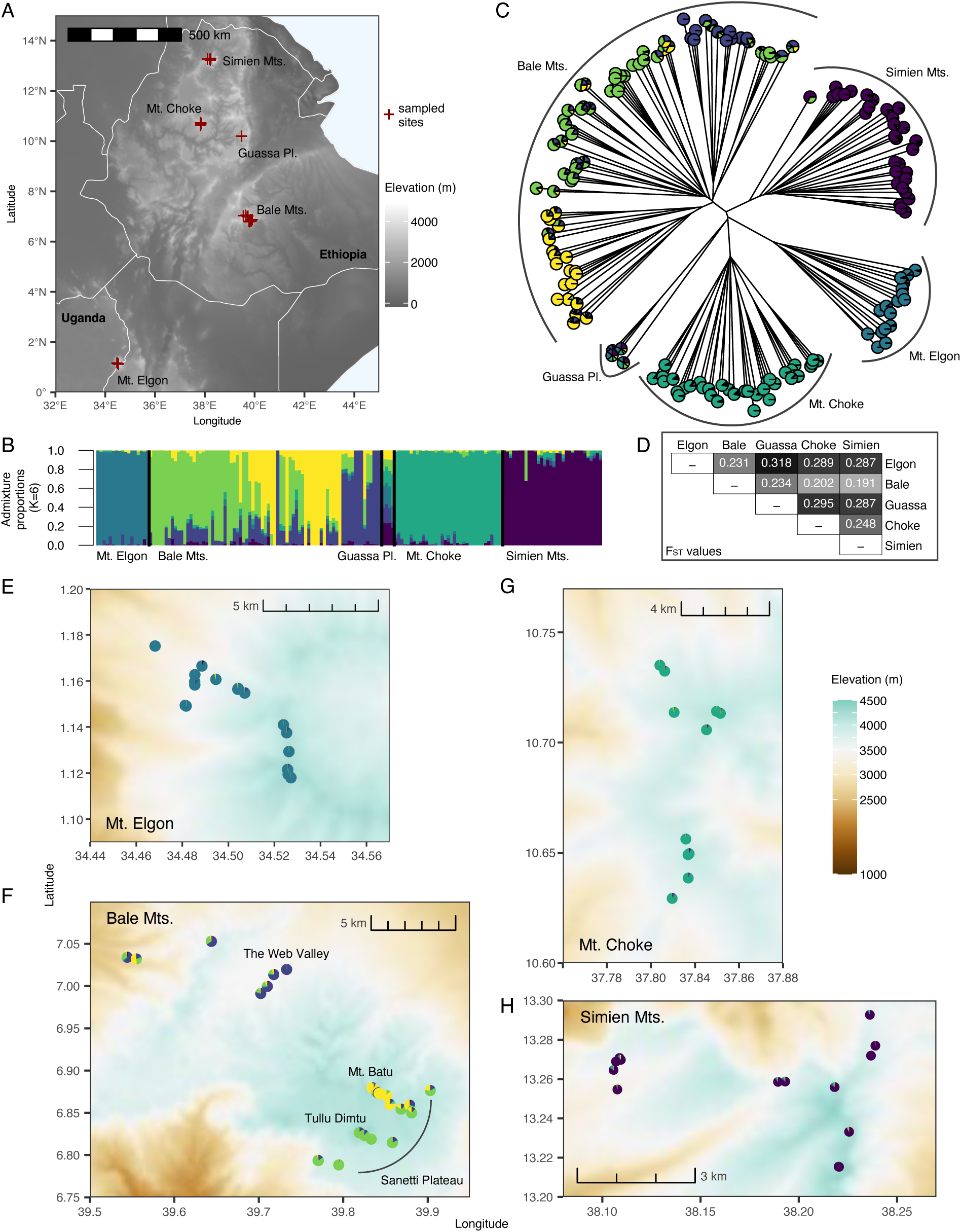
Sampling locations and population genetic structure of 163 Afroalpine genotypes using 146K unlinked SNPs. **A**. Map of study region showing sampled sites (red crosses) and elevation^49^. **B**. Genetic clustering per genotype based on sparse non-negative matrix factorization showing results for K=6. Individuals are sorted by latitude of origin. **C**. Neighbor-joining tree with tips colored by genetic clusters. **D**. F_ST_ between mountains. **E–H**. Zoomed-in sampled location in each mountain, showing elevational gradients and with individuals colored by genetic clusters.

Genome-wide differences between our studied Afroalpine populations suggest differences in adaptive potential (Supplementary Fig. 1). We quantified genome-wide nucleotide diversity (!), rarefied private allelic richness, and skew in the site-frequency spectrum (Tajima’s D). We found that ! (based on ∼2.6M sites with a mean depth of coverage of ∼35X across all samples) (Supplementary Fig. 1A) was greater in genotypes from the Bale Mts. (mean = 0.0014 ± 1.36e-06 se), followed by those from Mt. Elgon (mean = 0.0012 ± 1.19e-06 se), the Simien Mts. (mean = 0.0011 ± 1.27e-06 se), and Mt. Choke (0.0009 ± 1.06e-06 se). Rarefied private allelic richness followed the same pattern (based on ∼1.3M sites containing only per mountain private SNPs, Supplementary Fig. 1B): it was greater in genotypes from the Bale Mts. (mean = 1.26 alleles per site ± 0.0003 se), followed by those from Mt. Elgon (mean = 1.23 ± 0.0004 se), the Simien Mts. (mean = 1.15 ± 0.0003 se), and Mt. Choke (mean = 1.10 ± 0.0002 se).

Genome-wide Tajima’s D was generally close to 0 in all mountains (based on ∼2.6M sites, Supplementary Fig. 1C). Genotypes from the Bale Mts. had the greatest negative skew (mean = –0.52 ± 0.01 se), indicating more rare alleles, potentially capturing recent population expansion. Tajima’s D best reflected mutation-drift equilibrium in genotypes from the Simien Mts. and Mt. Elgon (mean ± se = –0.12 ± 0.01 and –0.03 ± 0.008, respectively). Genotypes from Mt. Choke had the greatest positive skew (mean = 0.22 ± 0.01 se), indicating more intermediate frequency alleles, potentially capturing recent population contraction or admixture, or due to limited sampling that might have not captured the breadth of genetic diversity.

Population size reconstructions of Afroalpine lineages indicate deep splits between mountains, congruent with the pattern of population genetic differentiation. Moreover, we found a shared trend of population decline towards the present (i.e., decreasing effective population size; N_e_), congruent with decreasing climatic suitability towards the end of the African Humid Period (Fig. 2). We inferred N_e_ history and divergence times using a framework that combines the site frequency spectrum and the recombination/linkage-disequilibrium information of a coalescent hidden Markov model^50^. Given that Arabidopsis thaliana is a highly selfing plant, we expected strong deviations from Hardy-Weinberg equilibrium. We thus maintain caution about interpreting the magnitude of Ne estimates. Furthermore, our germination experiments showed strong seed dormancy in many genotypes (results below), suggesting seed-banking, which could lead to a generation time greater than 1 year even though Arabidopsis is an annual. We thus use our results to interpret relative divergence times but suggest caution about whether they precisely identify time scales (*i.e.*, absolute time). To estimate historical habitat suitability, we first fit a species distribution model using contemporary species occurrences from East Africa and current Bioclim variables from the CHELSA v.2.1 climate database^51^. Bioclim variables included isothermality, temperature of the wettest quarter, precipitation of the driest month, and precipitation of the warmest quarter; chosen based on Pearson’s *r* < 0.7. We then projected this model into the past using the CHELSA-TraCE21k dataset^52^, which provides downscaled, high-resolution (1 km) transient climate data in 100-year steps from 21,000 years ago to the present. This allowed us to track the expansion and contraction of suitable habitat across 210 centennial time steps, accounting for post-glacial climate dynamics.

**Fig. 2.**
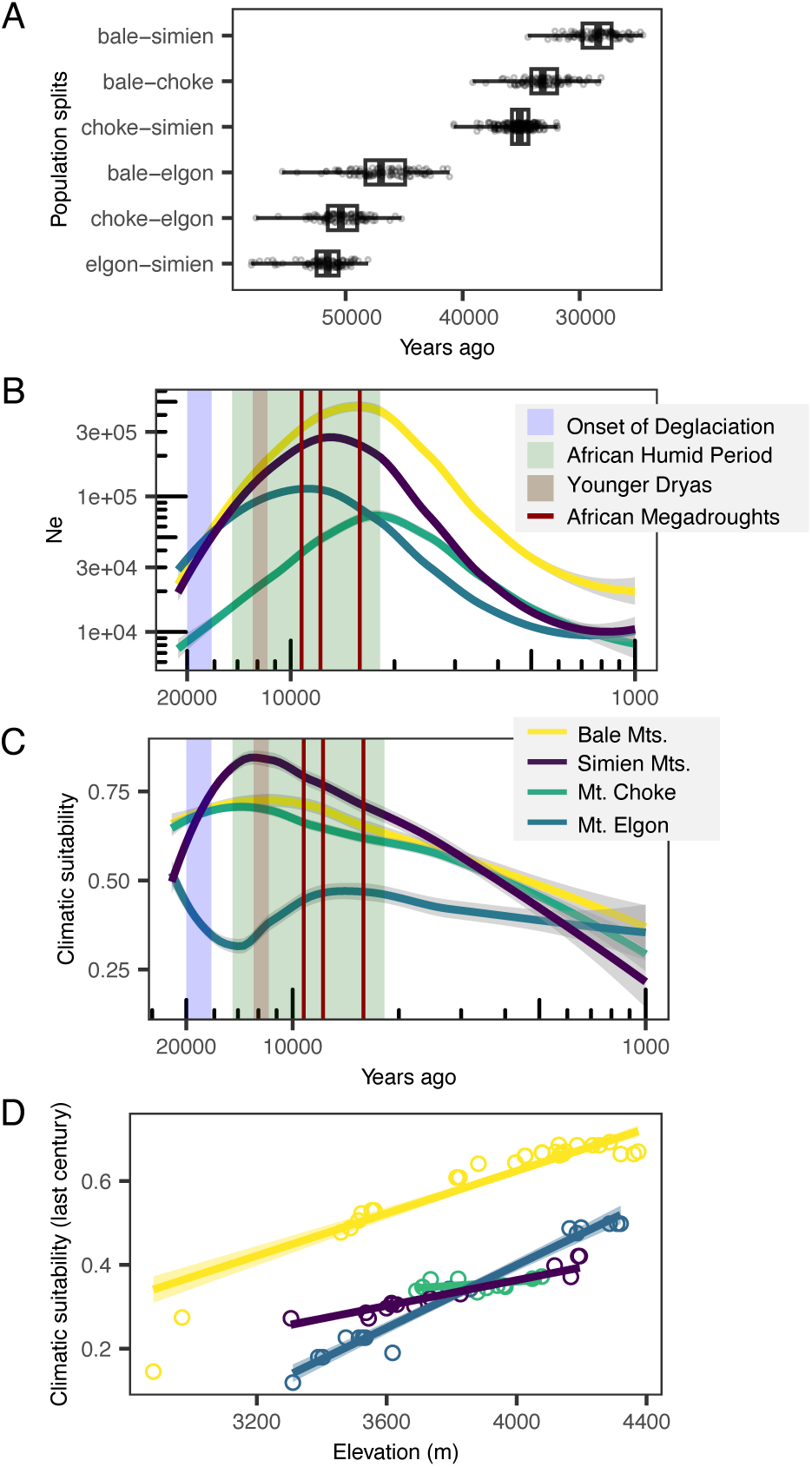
Genomic history and historical climatic suitability. **A**. Distribution of divergence times between mountain pairs estimated from cross-coalescence analysis (*n* = 100 random pairs). Whiskers of boxplots extend to extremes of split time estimates. **B**. Changes in *Ne* through time (here using 1.5 yr generation time). Bolt lines represent mean *Ne* per mountain over 100 bootstrapped joint models, surrounded by respective SE. **C**. Changes in climatic suitability through time. Bolt lines represent mean suitability across sampled sites at each mountain surrounded by respective SE. **D**. Climatic suitability, averaged across the last 100 years (from CHELSA-TraCE21k bioclim variables) at sampled sites (circles), increases with elevation and is highest in the Sanetti Plateau (Bale Mts.). Geological events in B and C were drawn based on the literature: Onset of deglaciation^47,48^, African Humid Period^54^, Younger Dryas^54^, African Megadroughts^54^.

Our results suggest a more recent divergence time (minimum ∼25–30K years ago) between the populations in the Ethiopian Highlands, and an older divergence time (minimum ∼45–50K years ago) among populations from East Africa more broadly (Fig. 2A). Our older dates (which still may be too recent because of seed banking) are comparable to the migration from East- into South-Africa and the expansion of the Eurasian clade, both dated at ∼45K years ago^44,53^ (using a 1 year generation time). Our estimated decrease in Ne towards the present might reflect a generalized worsening of environmental conditions for Arabidopsis across African sky-islands. When we use a mean generation time of 1.5 years (Fig. 2B), the timing of Ne changes better coincides with historical climatic events (Supplementary Fig. 2). We investigated the validity of this trend by estimating changes in climatic suitability since the LGM (Fig. 2C). We found that climatic suitability has sharply decreased starting ∼6.3K years ago, which coincides with the last severe drought that interrupted the African Humid Period (∼14.8–5.5K years ago) and marked the onset of increasing aridity in the region^54^. Over the last century, climatic suitability appears higher at higher elevations in all mountains and is overall higher in the Sanetti Plateau in Bale (Fig. 2D), highlighting the vulnerability of east African sky-islands refugia with climate change.

Despite striking similarities in habitats among mountains (Supplementary Fig. 3A–B), repeated evolution of alpine local adaptation in our studied African sky-island populations could be limited by heterogeneity of genomic histories, potential differences in climatic suitability, and strong climatic differences between mountains (Supplementary Fig. 3C–D). Below we first highlight inter-mountain differences in phenotypes from controlled experiments in growth chambers. Then we examine parallel local adaptation along elevation.

### Q2. Life history and plant size change with mountain of origin

Germination experiments revealed that genotypes from different mountains vary widely in seed dormancy and response to stratification treatments (Fig. 3A–B, Supplementary Fig. 4). We studied multidimensional seed dormancy in a subset of 66–74 genotypes by quantifying seed germination at different after-ripening times (0, 1, 3, and 6 months) combined with different stratification lengths (0, 7, and 30 days). Treatments were performed on seeds from mother plants grown under warm (22°C day/18°C night; 74 genotypes) and cool (12°C day/6°C night; 66 genotypes) conditions with a 12-hour photoperiod. Overall, genotypes from the northernmost and seasonal Simien Mts. showed strong dormancy and less variability among genotypes, while genotypes from the southernmost and humid Mt. Elgon showed weaker dormancy and more variability among genotypes (Supplementary Fig. 4). In all mountains, greater after-ripening times (≥ 3 months) stimulated germination, though germination was null or very low with no cold stratification.

**Fig. 3.**
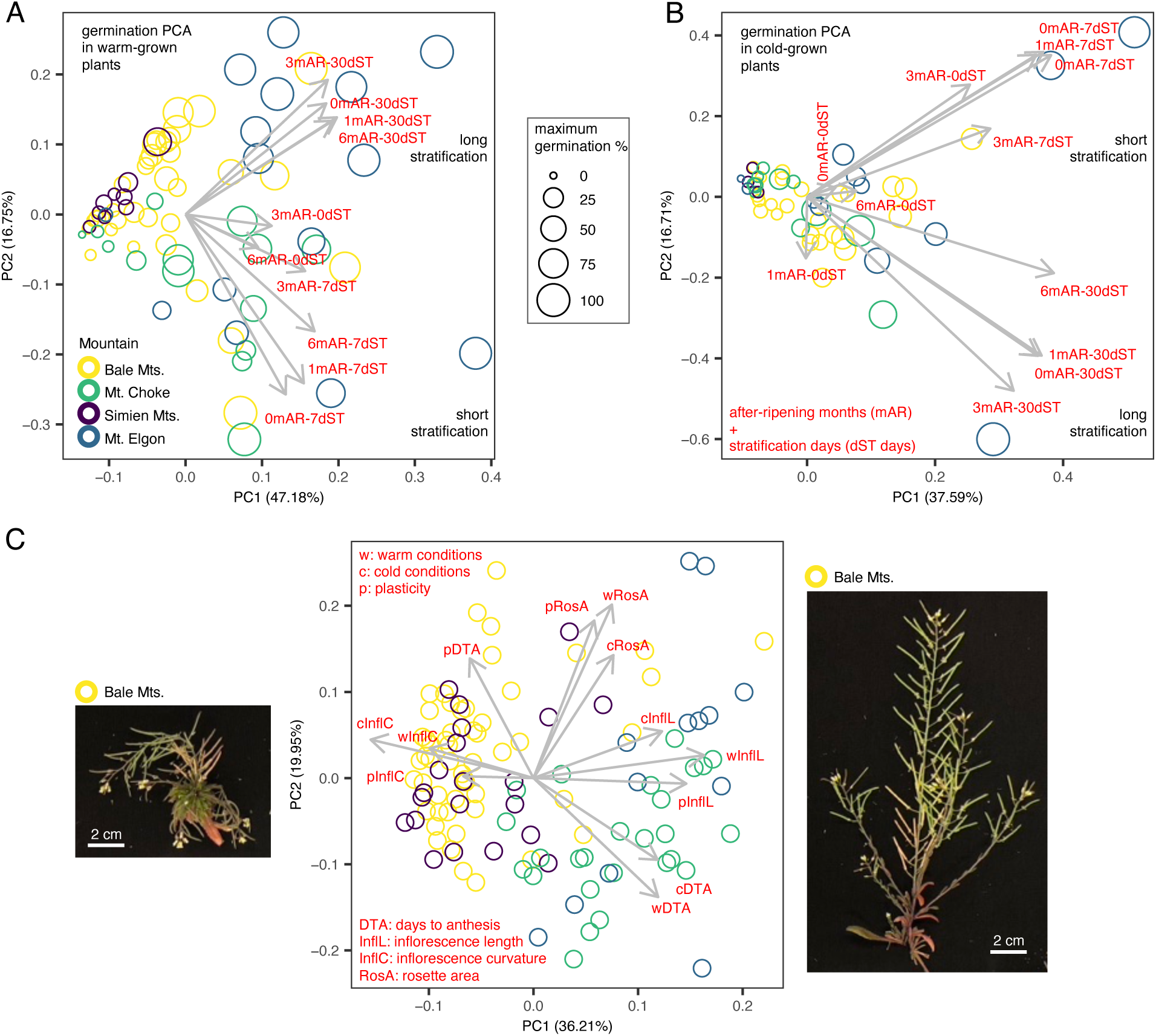
Principal component analyses of: **A**. Multidimensional seed germination phenotypes from plants grown in a warm maternal environment. **B**. Multidimensional seed germination phenotypes from plants grown in a cool maternal environment. **C**. Phenotypes from vegetative and reproductive plants in warm- and cold-growth chambers. Photos illustrate small and curved (left) and large and straight inflorescences (right) in warm-grown Bale genotypes. Genotypes are colored by mountains of origin.

To understand the potential coordinated multidimensional variation in seed germination behavior, we used principal component analysis (PCA). In seeds from warm-grown mother plants (Fig. 3A), PC1 (explaining 47% of the variation) was positively associated with greater germination across treatments, while PC2 (17%) was associated with germination response to stratification length: larger values representing greater germination with long stratification (30 days) and smaller values representing greater germination with short stratification (7 days). The segregation of genotypes along PC1 closely followed the strength of seed dormancy in each mountain; average maximum germination was lowest in genotypes from the Simien Mts. (mean ± se = 12 ± 5.5 %), followed by those from the Bale Mts. (mean ± se = 31 ± 4.9 %), Mt. Choke (mean ± se = 45 ± 9.5 %), and Mt. Elgon (mean ± se = 69 ± 7.8 %). In genotypes from the Simien Mts., full germination (>90% on a trial) was never reached; 64% was the maximum (6-month after-ripening and 30-day stratification treatment), in a genotype from 3305 masl, the lowest elevation sampled there. In Bale Mts. genotypes, germination mostly increased with longer stratification times (30-day; larger values in PC2), while in Mt. Choke genotypes, germination mostly increased with shorter stratification times (7-day; smaller values in PC2). In Mt. Elgon genotypes, germination response to stratification length was more diverse, with genotypes widely segregating along PC2. In seeds from cool-grown mother plants (Fig. 3B), which may better represent natural conditions for higher elevation plants, PC1 (38%) and PC2 (17%) followed a similar pattern as with warm grown mother plants described above, though overall germination was strikingly lower. It decreased on average by: ∼8X for the Simien Mts., ∼3X for the Bale Mts., and ∼2X for both Mt. Elgon and Mt. Choke.

Natural genetic variation of phenotypes from vegetative and reproductive plants in growth chambers was significant within and among mountains (Supplementary Figs. 5–6). We quantified trait variation in a subset of 113–131 genotypes by measuring flowering time (days to anthesis), inflorescence size, and rosette area, in plants grown under warm (22°C day/18°C night) and cold (8°C day/4°C night) conditions. Starting seeds came from the same warm-grown mother plants in germination experiments. In general, cold-grown plants had delayed flowering and decreased plant size compared to warm-grown ones (Supplementary Fig. 5). Phenotypic plasticity in response to cold (Supplementary Fig. 6) was similar between Mt. Choke and Mt. Elgon genotypes (higher plasticity in inflorescence length; greater shortening from warm to cold), and between the Bale and the Simien Mts. genotypes (higher plasticity in flowering time; greater delays from warm to cold).

To evaluate coordinated multidimensional variation in life history across warm and cold conditions, we used PCA (Fig. 3C). PC1 (explaining 36.2% of the variation) separated mountain pairs: most genotypes from the Simien and Bale Mts. grouped to the left (negative values); with short, curved inflorescences (*i.e.*, showing weak negative gravitropism), and rapid flowering, while most genotypes from Mt. Choke and Mt. Elgon grouped to the right (positive values); with long, erect inflorescences, and delayed flowering. No Mt. Elgon genotype had curved inflorescences, suggesting that curvature evolved in the Ethiopian Highlands. PC2 (20%) represented variation in rosette size, with genotypes from Mt. Elgon segregating most widely. In PC space, genotypes on the top left corner (mostly from the Bale Mts.) displayed larger rosettes and greater sensitivity to cold in flowering time (longer delays), while genotypes on the bottom right corner (mostly from Mt. Choke) displayed smaller rosettes and less sensitivity to cold in flowering time (shorter delays).

To examine how early life cycle traits (*i.e.*, seed germination behavior) might affect later life cycle traits (*i.e.*, flowering time and plant size), we used Pearson correlations. Covariation between seed germination and other phenotypes was sometimes consistent between mountains, or only present in single mountains. For instance, considering only warm-grown plants: in genotypes from the Simien Mts. and the Bale Mts. overall germination was higher (higher values of PC1 in Fig. 3A) in plants with longer inflorescences (Pearson *r* = 0.77 and 0.33, *n* = 9 and 31, respectively); while in genotypes from Mt. Choke and Mt. Elgon overall germination was higher in plants with shorter inflorescences (Pearson *r* = –0.55 and –0.45, *n* = 13 and 12, respectively). Moreover, in genotypes from Mt. Elgon long-stratification seed dormancy release (higher values of PC2 in Fig. 3A) occurred in plants with longer inflorescences (Pearson *r* = 0.61, n = 12). Lastly, in genotypes from the Simien Mts. plants with overall lower germination were rapid flowering (Pearson *r* = 0.62, n = 10). Taken together, the greater trait variation within Mt. Elgon and Mt. Choke suggests more potential for locally adaptive trait-elevation clines. We note, however, that our sampled environmental variability was limited in Mt. Choke (Supplementary Fig. 3C–D). In contrast, predominant strong seed dormancy, rapid flowering, and short/curved inflorescences within the Simien Mts. (and to some extent within the Bale Mts.) suggest bet-hedging that maintains low fitness (low fecundity) but allows escaping stressful conditions (*e.g.*, cold in the Bale Mts. and drought in the Simien Mts.).

### Q3. Parallel local adaptation along elevation is rare across mountains

Trait-elevation tests from linear mixed models that incorporated population genetic structure (*i.e.*, the allele similarity between genotypes representing the genome-wide pattern of variation) showed that putative local adaptation along elevation was rarely repeated across Afroalpine mountains, except for germination PC2 and inflorescence length (Fig. 4). When significant, these mixed models (below referred as ‘lmkin’) provide strong evidence of selection maintaining trait variation along an environmental gradient, because the relationship is independent from the genome-wide pattern of variation. Most trait-elevational clines were endemic, with stronger relationships in Mt. Elgon (Fig. 4C–H, J), weaker but significant in the Simien Mts. (Fig. 4A, C, G–I), while weak and mostly confounded with population genetic structure in the Bale Mts. (Fig. 4C, G–H). We found no evidence of trait-elevational clines in Mt. Choke, probably due to our limited elevational sampling there (∼380 m range), compared to other mountains (∼900 m in the Simien Mts., ∼1000 m in Mt. Elgon, and ∼1500 m in the Bale Mts.).

**Fig. 4.**
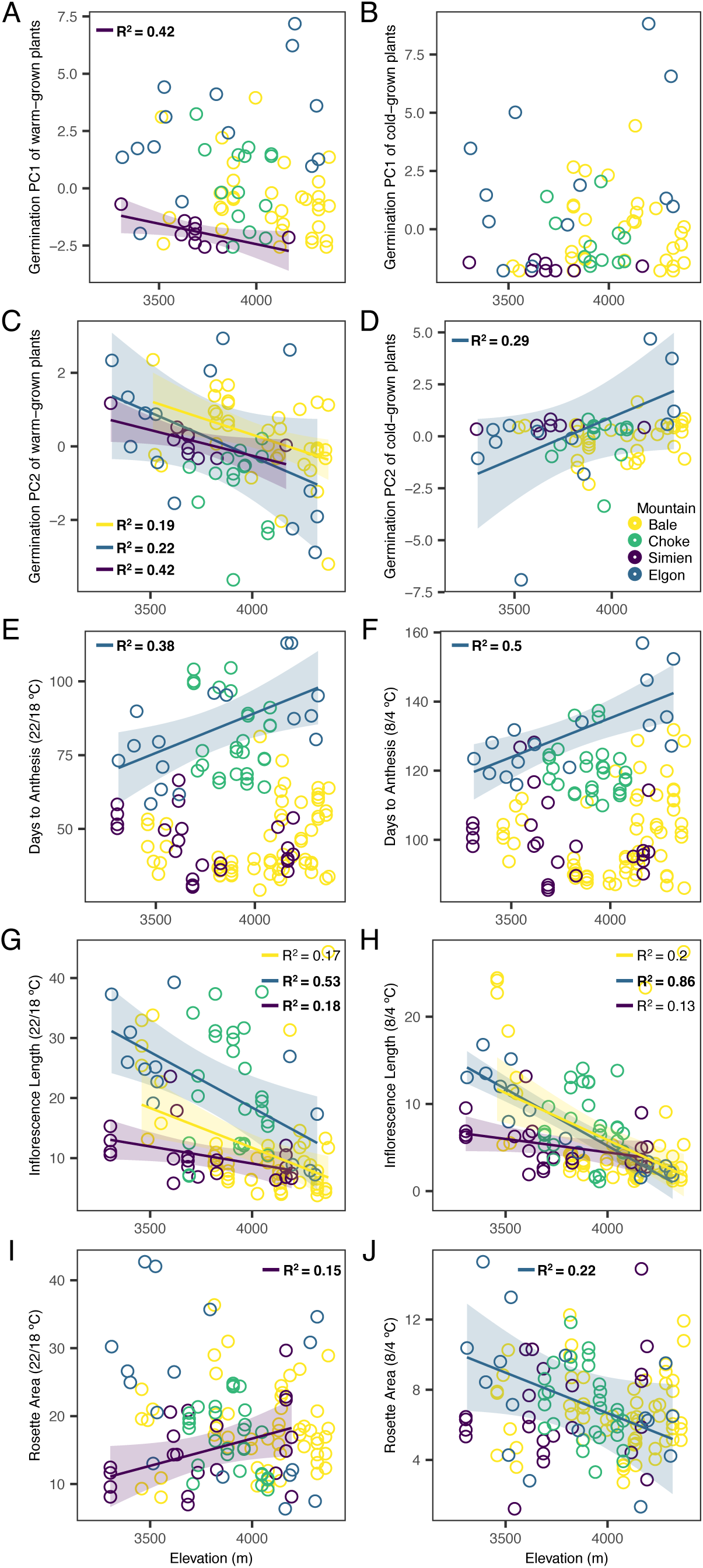
Mountain-specific trait-elevation clines of **A–D** germination PC1 and PC2, **E–F** days to flowering, **G–H** inflorescence length (in cm), and **I–J** rosette area (in cm^2^) measured in warm (left) and cold (right) growth chambers. Linear relationships (± se) and R^2^ come from standard linear regressions that were significant (*P* ≤ 0.05). The R^2^ is bolded if relationships were significant in linear mixed models that accounted for kinship between genotypes (*P*_lmkin_ ≤ 0.05). Genotypes are colored by mountains of origin.

Despite contrasting climates and strong phenotypic differences, genotypes from Mt. Elgon and the Simien Mts. displayed evidence of elevational selection (*P*_lmkin_ ≤ 0.05) in seed dormancy-release via stratification length and inflorescence length in warm-grown plants (Fig. 4C, G). Sky-islands genotypes (*i.e.*, those at the highest elevations) shared a phenotype of seed-dormancy release after short (7-day) stratification (lower values of germination PC2; Mt. Elgon *P*_lmkin_ = 0.04, *n* = 14, Simien Mts. *P*_lmkin_ = 0.007, *n* = 10) and short inflorescences (Mt. Elgon *P*_lmkin_ = 1.3×10^−4^, *n* = 13, Simien Mts. *P*_lmkin_ = 0.03, *n* = 21). These trait-elevation clines were also significant in cold-grown plants, but only in Mt. Elgon genotypes (Fig. 3D, H; germination PC2 *P*_lmkin_ = 0.03, *n* = 12, inflorescence length *P*_lmkin_ = 2.5×10^−7^, *n* = 16). Sky-island genotypes from the Bale Mts. showed similar patterns (Fig. 4C, G–H), as well as more prostate (curved) inflorescences under cold conditions (Supplementary Fig. 7B), but evidence of selection was only seen for germination PC2 (*P*_lmkin_ = 0.004, *n* = 35). Sky-islands genotypes in Mt. Elgon also had significantly delayed flowering in both warm-grown and cold-grown plants (Fig. 4E–F, *P*_lmkin_ = 0.002, *n* = 16; *P*_lmkin_ = 6.6×10^−5^, *n* = 16, respectively), perhaps similar to mountainous Mediterranean winter annuals. These genotypes also had smaller rosettes, though only when grown under cold conditions (Fig. 4J, *P*_lmkin_ = 0.03, *n* = 16). In contrast, sky-islands genotypes in the Simien Mts. had the strongest seed dormancy (lower values of germination PC1; Fig. 4A, *P*_lmkin_ = 0.007, *n* = 10), consistent with bet-hedging and potentially seed-banking, perhaps similar to lowland Mediterranean rapid cyclers. These genotypes also had larger rosettes (*P*_lmkin_ = 0.025, *n* = 21), though only when grown under warm conditions (Fig. 4I).

### Q4. High-elevation genotypes are rarely more sensitive to cold than low-elevation ones

In general, high-elevation genotypes did not differ in cold sensitivity (*i.e.*, response to cold) compared to low-elevation ones, except in the Simien Mts. We quantified plasticity/sensitivity in response to cold as the change in magnitude of trait values measured under cold relative to warm conditions (*i.e.*, trait value under warm minus under cold). Sensitivity to cold was higher in the high-elevation Simien Mts. genotypes for rosette size (Fig. 5C); *i.e.*, genotypes from higher elevations produced significantly larger rosettes when grown under warm compared to cold conditions (*P*_lmkin_ = 0.01, *n* = 21). Producing larger rosettes under warm conditions could indicate that high-elevation Simien Mts. populations change their rosette size in years when conditions are more favorable.

**Fig. 5.**
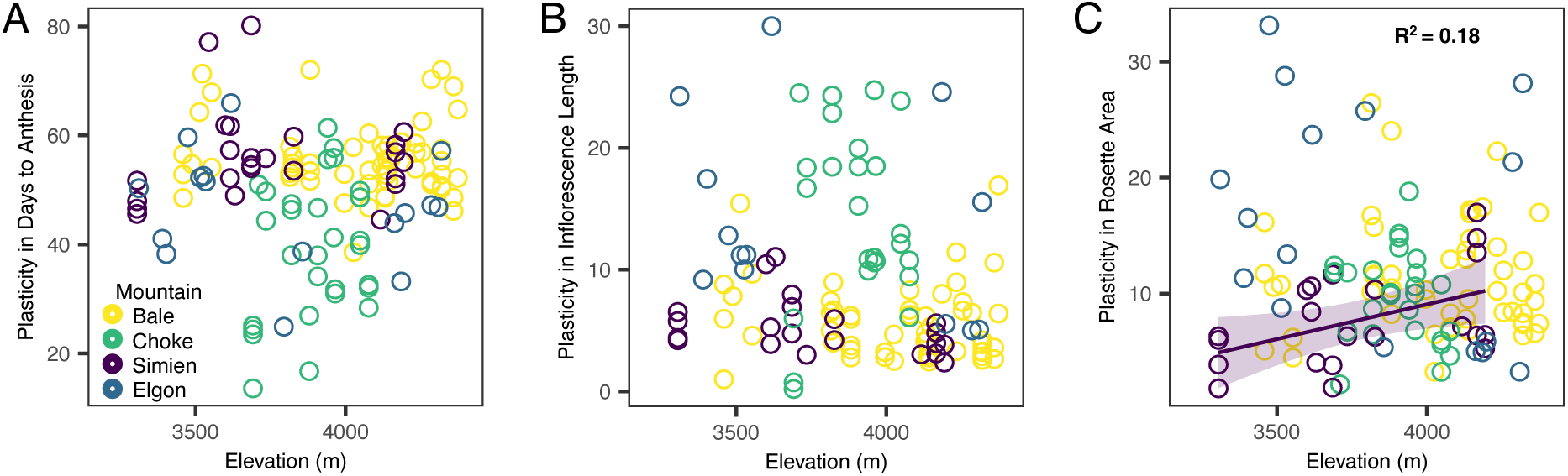
Mountain-specific change in trait cold-response with elevation of origin in **A**. Flowering time, **B**. Inflorescence length (in cm), and **C**. Rosette Area (in cm^2^). Linear relationships (± se) and R^2^ come from standard linear regressions that were significant (*p* ≤ 0.05). The R^2^ is bolded if relationships were significant in linear mixed models that accounted for kinship between genotypes. Genotypes are colored by mountains of origin.

### Q5. Genetic similarity likely shapes allele reuse in quantitative trait loci

Our GWAS results revealed putative quantitative trait loci (QTL) of inflorescence length and days to anthesis (in warm-grown plants) composed each of one SNP located upstream (in the putative promoter region) and downstream (in the putative enhancer region), respectively, of *TEOSINTE BRANCHED 1*/*CYCLOIDEA*/*PCF5* (*TCP5*) (Fig. 6A). In *Arabidopsis thaliana*, this transcription factor positively regulates plant thermomorphogenesis^55^ and overexpression causes short, highly branched and bushy inflorescences, and sterile pollen^56^. We implemented GWAS using univariate mixed models that accounted for population genetic structure in a random subset of 78 genotypes that excluded quasi-clones (based on genetic dissimilarity < 0.5).

**Fig. 6.**
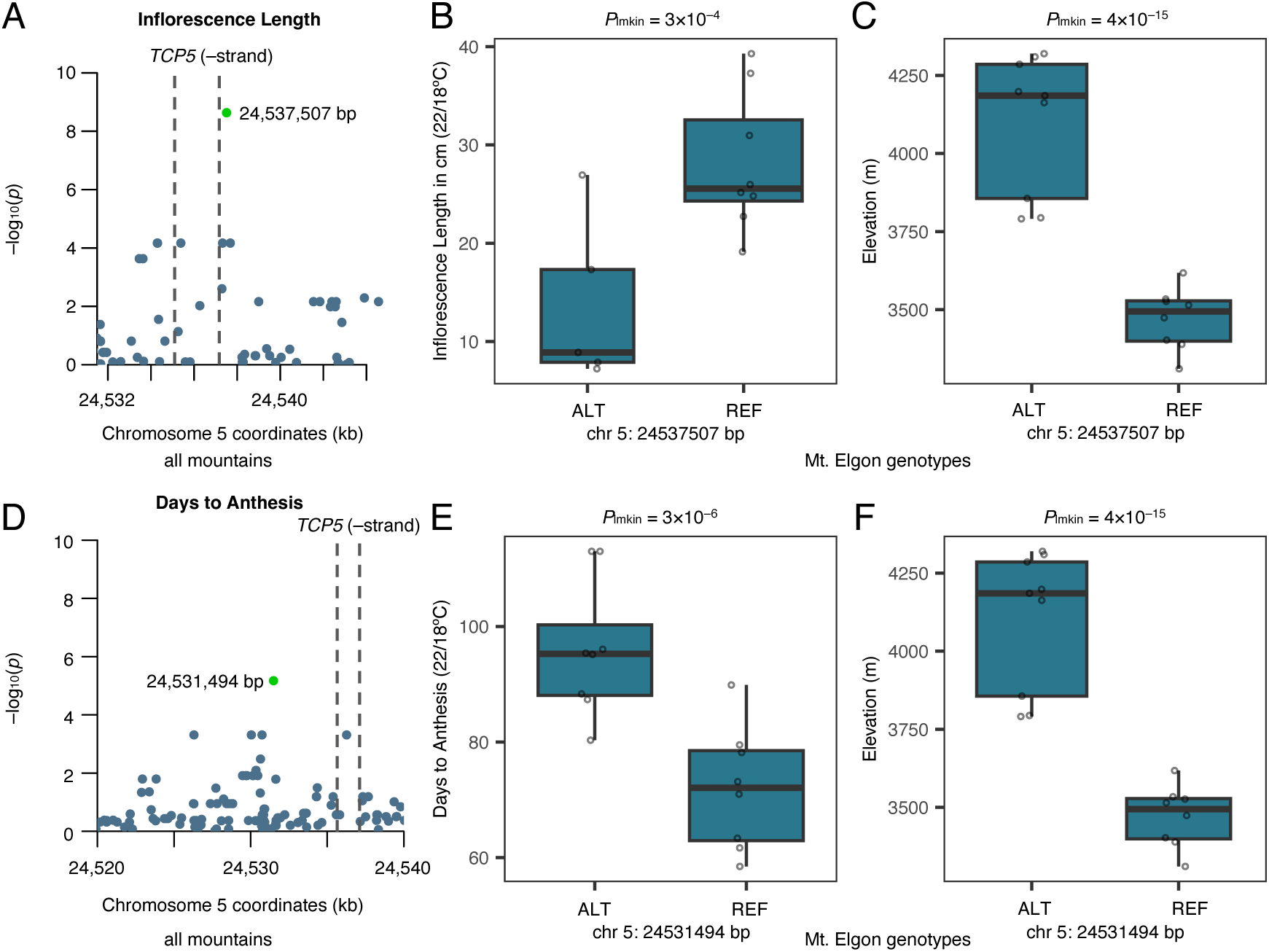
Quantitative trait locus for inflorescence length and flowering time mapped in studied East African genotypes with genome-wide association studies. **A, D**. Zoomed-in Manhattan plots showing top SNPs in green (FDR=0.002 and 0.12, respectively). **B, E**. Effect size in Mt. Elgon genotypes (MAF=0.5). **C, F**. Elevational variation of top SNP in Mt. Elgon genotypes.

For inflorescence length in warm-grown plants, we analyzed 66 genotypes and 701,984 SNPs (Supplementary Fig. 8A). The top SNP (chromosome 5: 24537507 bp; Fig. 6A) had an allele frequency (AF) of 0.258 (*P* = 2.3×10^−9^, FDR = 0.002), but only segregated in genotypes from Mt. Elgon (AF = 0.5) and Mt. Choke (AF = 0.241), and only in Mt. Elgon allele frequency changed with elevation (*P*_lmkin_ = 4.4×10^−15^, *n* = 17). In Mt. Elgon, the short-inflorescence allele was more frequent in genotypes from higher elevations (Fig. 6B–C). The Bale and Simien Mts. genotypes were fixed for the short-inflorescence allele, though GWAS might have not identified the causal SNP altering inflorescence size.

For flowering time (days to anthesis) in warm-grown plants, we analyzed 76 genotypes and 755,492 SNPs (Supplementary Fig. 8B). The 2^nd^ top SNP (chromosome 5: 24531494 bp; Fig. 6D) had an AF of 0.724 (*P* = 6.7×10^−6^, FDR = 0.12), but only segregated in genotypes from Mt. Elgon (AF = 0.5; in linkage with chromosome 5: 24537507 bp), where allele frequency changed with elevation (*P*_lmkin_ = 4.4×10^−15^, *n* = 17). The late-flowering allele was more frequent in genotypes from higher elevations (Fig. 6E–F). The Bale and Simien Mts. genotypes were fixed for the early-flowering allele, while Mt. Choke genotypes were fixed for the late-flowering allele.

We found that allele reuse was greatest between genetically similar mountains (*i.e.*, those in the Ethiopian Highlands), and lowest between Mt. Elgon and the rest (Supplementary Table 1). To evaluate allele reuse, we calculated Pearson correlations between mountain pairs’ minor allele frequencies obtained from GWAS SNPs with the 100-top *P*-values for flowering time, inflorescence length, rosette size, and the germination PC1 and PC2 in warm-grown plants (phenotypes with greater variation). Genotypes from the Simien and the Bale Mts. showed the greatest allele reuse (*r* > 0.96, *P* < 2.2×10^−16^ for all phenotypes). Moreover, germination PC2 showed the greatest allele reuse across all mountain pairs (*r* ≥ 0.6, *P* ≤ 3×10^−16^), though correlations were greater between Ethiopian mountain pairs (*r* > 0.9, *P* = 2×10^−16^). Although genotypes from Mt. Elgon and Mt. Choke were phenotypically similar (Fig. 3C), allele reuse was only apparent for SNPs associated with flowering time (*r* = 0.72, *P* < 2×10^−16^). Thus, allele reuse appears limited between mountains that are genetically divergent despite being phenotypically similar (*i.e.*, Mt. Choke and Mt. Elgon; Fig. 1D). However, allele reuse was significant between genotypes from Mt. Elgon and the Simien Mts. for inflorescence length and germination PC2 (*r* = 0.7, *P* = 2×10^−16^, and *r* = 0.6, *P* = 3×10^−16^), traits that showed evidence of repeated elevational selection (Fig. 4C, G).

## Discussion

Local adaptation along elevation is a key driver of intraspecific diversity in animals and plants, and because mountains are often isolated, these systems are a model for studying the predictability of evolution^57^. The model plant *Arabidopsis thaliana* displays significant natural genetic variation along elevation in ecologically important traits, suggesting locally adaptive elevational clines in different parts of its native range^33,36–38^. Putative locally adaptive elevational clines, however, can shift between populations on opposite sides of Eurasia, indicating that increasing geographic isolation, as well as genetic and climatic heterogeneity between mountains, reduces the probability of parallel alpine adaptation^35^. For example, delayed flowering and high-water use efficiency appear adaptive at high elevations in the western Mediterranean, consistent with an alpine conservative/slow strategy. However, early flowering and low water use efficiency appear adaptive at high elevations in Asia, consistent with an alpine acquisitive/rapid-cycling strategy^35^. While elevational adaptation has been relatively well-studied in the Eurasian range of *Arabidopsis thaliana*, the older, long isolated, and potentially threatened, sky-island populations of East Africa remain little known^42–45^. Here we examined parallel adaptation to high elevation in *Arabidopsis thaliana* populations from four isolated mountains in the Ethiopian Highlands and a volcano in Uganda. We combined whole genomes and growth chamber experiments to study genomic history and natural genetic variation in life history traits. We tested whether different mountain peaks share a common Afroalpine strategy with a shared genetic basis.

### Life history strategies in tropical alpine ecosystems differ between mountains

We found that mountain populations have long been genetically isolated, potentially differ in adaptive potential, and have strong phenotypic differences. While mountains share a trend of decreasing effective population size and climatic suitability towards the present, both genetic diversity and climatic suitability differed between mountains (Fig. 2, Supplementary Fig. 1). Moreover, we uncovered strong inter-mountain differences in ecologically important traits, and such differences were not clearly geographically structured. Genotypes from the Simien and the Bale Mts., on opposite sides of the East African Rift Valley, were more similar to each other, while genotypes from Mt. Choke and Mt. Elgon, separated by considerable geographic distance (∼900 km), were more similar to each other (Fig. 3). Accordingly, we found limited evidence of parallel Afroalpine adaptation along elevation. Putatively adaptive elevational clines shared between mountains were evident for only two traits (shorter inflorescences and seed dormancy release with short stratification at higher elevations; Fig. 4), but with potentially different genetic bases. Surprisingly, these clines were strongest in genotypes from the two most genetically divergent, climatically distinct, and phenotypically different mountains: the northernmost seasonally dry Simien Mts. and the equatorial more humid Mt. Elgon. Evidence of parallel elevational adaptation in the rest of traits was absent; trait (and trait plasticity)-elevational clines were either endemic to single mountains or confounded with population genetic structure. Our results indicate that East African sky-islands harbor different alpine strategies where each mountain top is a reservoir of distinct genetic diversity in traits. Thus, instead of a “global” tropical alpine strategy across East Africa^46^, we find rather “endemic” tropical alpine strategies.

Phenotypic differences between mountains were mediated by both life history and plant size traits. Genotypes in the climatically divergent Simien Mts. displayed strong seed dormancy, early flowering (*i.e.*, stress escape), and very short inflorescences, under both warm and cold conditions. This is consistent with a risk-avoidance bet-hedging strategy, which was more pronounced at higher elevations: plants in unpredictable, harsh environments reduce year-to-year fitness variation by sacrificing short-term maximum yield^22^. Strong seed dormancy promotes seed banking, allowing long-term persistence; if a harsh frost or drought kills emerging seedlings, ungerminated seeds serve as an evolutionary safety net, allowing the population to persist and germinate in a subsequent, more favorable year^21^. Genotypes from the somewhat colder Bale Mts. also displayed bet-hedging (strong seed dormancy and early flowering), but diversity in traits was much higher there compared to the Simien Mts (Supplementary Fig. 4–6). Accordingly, genetic diversity and climatic suitability was highest in the Bale Mts. (Supplementary Fig. 1, Fig. 2), likely because of its wide extent of Afroalpine habitat in the region. For example, the Bale Mts. are the center of genetic diversity in the Afroalpine species *Trifolium cryptopodium* and *Carduus schimperi*^58^. Genotypes in the less seasonal and relatively more humid Mt. Elgon displayed the greatest diversity in all traits and the strongest trait-elevation clines, showing a coordinated delayed flowering and seed dormancy release with short stratification associated with shorter and seemingly bushier inflorescences at higher elevations (Fig. 4). Thus, selection along elevation is likely more important in maintaining trait diversity in Mt. Elgon than in other mountains, somewhat similar to mountainous Iberian populations^32^. Genotypes in Mt. Choke were surprisingly more phenotypically similar and as variable as those in Mt. Elgon, despite our more limited sampling there.

### Genetic convergence in quantitative trait loci decreased with genetic divergence

We found that allele reuse in quantitative trait loci mapped with GWAS was higher between more closely related mountains, particularly between the phenotypically similar Simien and Bale Mts. Although Mt. Elgon genotypes were more phenotypically similar to Mt. Choke genotypes, evidence of allele reuse was almost absent (except for flowering time). Recent evidence suggests that the probability of genetic convergence (here defined as similar phenotypes with shared genetic bases) in alpine adaptation in the genus *Arabidopsis* is influenced by divergence time between study lineages^10^. Genetic convergence is predicted to be stronger between closely related populations or species^11^, although pervasive gene co-option for similar ecological strategies across deep timescales is also possible^8^. Within species, it thus seems likely that genetic convergence is common, but our previous study showed that genetic and environmental heterogeneity in natural populations of mountainous Eurasian *Arabidopsis thaliana* limit evolution of repeated local adaptation along elevation^35^. Although here we studied more geographically proximate mountain populations than in Eurasia, the relative isolation was likely greater due to the unsuitability of lowland climates in East Africa for Arabidopsis. We also find striking ecological differences, along with deep genetic divergence, despite populations facing a similar warm/bright day-freezing night challenge. Because one mechanism of genetic convergence depends on isolated populations having shared standing genetic variation, it is likely that the disjunct nature of high-altitude sky-islands in East Africa limits gene flow, promoting localized independent evolution with different genetic bases. For instance, we found that high-altitude dwarf inflorescences and delayed flowering might be associated with variation in the *TCP5* transcription factor in Mt. Elgon (Fig. 6), but not in the Ethiopian Highlands.

Afroalpine tropical sky-islands are hotspots of biodiversity and endemism^4,5,15^. These isolated mountain ranges and peaks surrounded by vast, inhospitable lowlands, are however critically endangered ecosystems due to climate change^5,15^. By measuring traits associated with life history (seed dormancy, flowering time, plant size), integrated with genomic and climatic history reconstructions, we uncovered the diversity of tropical Afroalpine strategies in the model plant *Arabidopsis thaliana*. Our findings highlight that predicting evolution under seemingly similar ecological challenges is complicated by deep eco-evolutionary heterogeneity. This is unlike scenarios of more recent colonization events to novel environments, where adaptive mechanisms seem more predictable (*e.g.*, Cape Verde colonization of *Arabidopsis thaliana* ∼5–7K years ago^59^). Sky-islands in East Africa are rather “cradles” of diversity in traits, thus important for preserving the full adaptive potential of *Arabidopsis thaliana*, and in general of the iconic Afroalpine flora.

## Methods

### Plant material

In 2017–2018, we collected fruiting plants in Ethiopia and Uganda covering elevational gradients of ∼380–1500 m. Samples included the Bale Mts.: 75 individuals in 29 sites from 2881– 4374 m asl; Mt. Choke: 35 individuals in 11 sites from 3691– 4077; the Simien Mts.: 32 individuals in 14 sites from 3305– 4194, and Mt. Elgon: 17 individuals in 17 sites from 3311– 4320 m. We also included four plants from the Guassa Plateau collected in two sites at 3426 and 3450 m asl. To increase seed and obtain tissue for DNA, field collected seeds were stratified for seven days in 1000 ppm gibberellic acid (GA) at 4°C, sown in potting mix, grown in a growth chamber (24°C day, 20°C night, with a 16-hour photoperiod, 150-μmols m s−1 light), and kept well-watered at the Pennsylvania State University. DNA plants were harvested vegetatively after 4–6 weeks and preserved in silica gel until processing. Seed-increase plants were thinned to one plant per pot and transferred after two weeks of germination to a dark cold chamber (4°C) for a 6-week vernalization. Plants were transferred back to the original growth chamber (24°C day, 20°C night, with a 16-hour photoperiod) and kept well-watered until harvesting. Here and in subsequent experiments, trays were periodically rotated (twice per week) and moved around the growth chamber (twice per month) to mitigate positional effects. Seeds were harvested when plants had >80% ripe siliques and kept in dry-cold conditions until conducting growth chamber experiments.

### Climate data

We downloaded global gridded data for bioclimatic variables at 30 arcsec (∼1km^2^) spatial resolution from 1981 until 2010 included in the CHELSA-BIOCLIM+ dataset v.2.1^51^. We extracted bioclimatic data based on coordinates at collection sites in R (R Core Team, 2014). Coordinates were transformed to spatial points in the WGS84 Coordinate System (same as bioclimatic data) with the SpatialPoints function in the R package sp (v.2.2-1)^60^, and bioclimatic data was obtained using the extract function in the R package terra (v.1.9-25)^61^. The final environmental dataset included 48 variables. Environmental gradients of our study region were identified with PCA (R function prcomp, variables scaled and centered) (Supplementary Fig. 3C–D). Elevation of origin was recorded directly in the field with a GPS.

### Sequencing data

Fifteen genotypes with provenance data were previously sequenced^35,42^. For the other 148 genotypes, total genomic DNA was extracted from ∼100 mg of fresh tissue collected from growth chamber individuals, using the Viogene Plant Genomic DNA Extraction System (Viogene BioTek, Corp.). To ensure purity and high molecular weight, DNA was subsequently cleaned with 0.9X AMPure XP magnetic beads (Beckman Coulter, Inc.). DNA purity was assessed with a Thermo Scientific NanoDrop 2000 spectrophotometer (Thermo Fisher Scientific, Inc.) and quantified with an Invitrogen Qubit 2.0 fluorometer (Thermo Fisher Scientific, Inc.). Individuals were sequenced at an average coverage of ∼50X per genotype at the Texas A&M AgriLife Research (Genomics and Bioinformatics Service). At least 500 ng of genomic DNA was used to construct paired-end sequencing libraries (PerkinElmer NEXTFLEX Rapid XP DNA-Seq Kit HT) which were sequenced on an Illumina NovaSeq X 25B Single Lane platform (Illumina, Inc.). Sequence cluster identification, quality prefiltering, base calling and uncertainty assessment were done in real time using Illumina’s NCS 1.0.2 and RFV 1.0.2 software with default parameter settings. Sequencer cbcl basecall files were demultiplexed and formatted into fastq files using bcl2fastq 2 2.19.0 script configureBclToFastq.pl. Raw reads were processed with FastQC (v.0.11.8)^62^ and filtered for low quality and adapter regions using Trimmomatic (v.0.39)^63^. Filtered files contained on average 30M reads per genotype.

### SNP genotyping

Paired-end reads from the 163 genotypes were mapped to the *Arabidopsis thaliana* TAIR10 reference genome using the Araport11 reannotation^64^ using BWA-MEM (v.0.7.15)^65^ with default parameter settings. Mapped reads were processed, and SNP detection was performed, as in Gamba et al. (2024)^35^, following GATK v.4.2.6.1 best practices for obtaining biallelic SNPs using hard filters to ensure high quality variants^66^. We then used Beagle v.5.2^67^, to phase genotypes and impute missing SNPs based on linkage disequilibrium with default parameter settings. To remove potential paralogs, sites with excess heterozygosity (flagged ‘ExcHet<1’) and with >5% heterozygote genotypes were filtered out. The final set of 2,611,238 high-confidence biallelic SNPs had a mean mapped depth of coverage of 35.4X across all samples (range: 15.2X–90.5X).

### Seed germination phenotyping

Seeds were stratified in 1000 ppm in GA for seven days, sown in potting mix, and thinned to one plant per pot, using eight replicates (pots) per genotype. Plants were germinated in a growth chamber (24°C day, 20°C night, with a 16-hour photoperiod, 150-μmols m s−1 light) and after 2 weeks, seedlings were transferred to a dark cold chamber (4°C) for a 6-week vernalization. Plants were then transferred to a “warm” growth chamber (22°C day, 18°C night, 12-hour photoperiod, 200-μmols m s−1 light) until bolting. To test for the effect of seed maturation environment on germination, we kept four replicates in the “warm” growth chamber and transferred four replicates to a “cool” growth chamber (12°C day, 6°C night, 12-hour photoperiod, 200-μmols m s−1 light, simulating high-elevation conditions) as soon as plants began bolting. Seeds from both “warm” and “cool” chambers were harvested when > 80% of the siliques were ripe. The final number of genotypes used for germination trials below were 74 from the “warm” maturation environment and 66 from the “cool” maturation environment, based on the number of seeds and space available, and covering the elevational gradients of sampled mountains.

To determine what after-ripening and stratification times promote germination/break dormancy, fresh seeds from each maturation chamber were sorted into 12 germination treatments. Harvesting in the “cool” maturation chamber started ∼2 months after harvesting in the “warm” maturation chamber. Depending on harvesting time of replicates, we performed two simultaneous or consecutive trials per genotype per seed germination treatment. Treatments included four after-ripening times under dry and dark conditions at room temperature (10-days/0-month, 1-month, 3-months, and 6-months), followed by three stratification times under dark and cold conditions (4°C, simulating high elevation) in 2 ml Eppendorf’s filled with DI water (0-day, 7-days, and 30-days). To score germination, we used 24 seeds per trial per genotype, arranged in a 6 ’ 4 grid in a 100 x 15 mm polystyrene petri dish (VWR International), lined with blue germination paper (Hoffman Manufacturing). To prevent fungal growth, seeds were imbibed in 35 ml of a 2% PPM solution (Plant Preservative Mixture, Caisson Laboratories). Petri dishes were placed in a growth chamber under germination-inducing conditions at 22 °C and 12-hour photoperiod (100-μmols m s−1 light). Total germination was manually scored based on radicle protrusion after 10 days in a total of 3,360 petri dishes. Germination percent was averaged between trials to obtain one value per genotype per treatment.

### Growth chamber phenotyping

Starting seeds came from the same plants in the “warm” maturation environment above, stratified in 1000 ppm in GA for seven days, sown in potting mix, and thinned to one plant per pot, using six replicates (pots) per genotype. Plants were germinated in a growth chamber at constant temperature (24°C, 13-hour photoperiod, 150-μmols m s−1 light) and after 2 weeks, seedlings were transferred to a dark cold chamber (4°C) for a 6-week vernalization. Three replicates were then placed in a “warm” growth chamber (22°C day, 18°C night, 12-hour, photoperiod, 200-μmols m s−1 light) and three replicates in a “cold” growth chamber (8°C day, 4°C night, 200-μmols m s−1 light). To track plant growth, trays (containing 31 pots each) were photographed at a constant distance with a ruler weekly until bolting. To track flowering phenology, plants were monitored every other day. For measuring inflorescence length, inflorescences with >75% ripe siliques were cut with scissors from the base, placed on a black cloth, and photographed at a constant distance with a ruler.

We measured phenotypes on up 114–131 genotypes with 2–3 replicates that survived until harvesting, for a total of 113 genotypes in both “warm” and “cold” experiments. Phenotypes included: final rosette area measured in cm^2^ from photographs of bolting rosettes, days to anthesis here defined as days until white petals appeared, inflorescence length measured in cm from photographs, and inflorescence curvature scored as 1 (present) or 0 (absent) from photographs. Measurements from photographs were performed with ImageJ^68^. After quality/error checking, the best linear unbiased estimate (BLUE) of phenotypes was calculated per genotype with the BLUE function in the R package polyqtlR (v.0.1.1)^69^, using genotype as the predictor of trait measurements across replicates and tray as a random effect.

### Population genetic structure

We used two clustering methods to infer population genetic structure that we interpret collectively. First, we thinned our dataset of 2,611,238 SNPs to exclude variants in high linkage-disequilibrium with the snpgdsLDpruning function in the R package SNPRelate (v.0.9.19)^70^, pruning SNPs with correlation > 0.5 in 100 kb sliding windows. The resulting dataset contained 146,431 unlinked SNPs. Then, we calculated admixture proportions for each genotype. Because Hardy-Weinberg equilibrium is likely violated in a selfer like *Arabidopsis thaliana*, we calculated admixture proportions based on sparse nonnegative matrix factorization algorithms in sNMF^71^, implemented in the R package LEA^72^, using a regularization parameter of alpha=100. Cross-validation of the number of clusters was determined from log-likelihoods of the sNMF output across 20 replicates for K=1–15, supporting K=6. Subsequently, we computed an unrooted phylogenetic tree with the Neighbor-Joining (NJ) algorithm^73^ based on a genetic dissimilarity matrix using all unlinked SNPs. We inferred genetic dissimilarity with the snpgdsDiss function in SNPRelate. The tree was plotted with the plotnj function in the R package phyclust (v.0.1-33)^74^ and tips were colored according to their admixture proportions using the tiplabels function and the pies option in the R package ape (v.5.7-1)^75^. Finally, we calculated Nei’s^76^ pairwise F_ST_ with function pairwise.fst.dosage in the R package hierfstat (v.0.5-11)^77^. For this, a genotype matrix with dosage data for unlinked sites, excluding SNPs with MAF < 0.05, was estimated with the function snpgdsGetGeno in the R package SNPRelate.

### Genome-wide diversity

We calculated nucleotide diversity !^78^ for each mountain using our dataset of 2,611,238 sites. Per mountain vcf datasets containing all sites were generated with BCFtools (v.1.18)^79^ (view -S), and genome-wide estimates of ! were obtained with VCFtools (v.0.1.15)^80^ using 50 kb sampling windows. Deviations from neutral evolution in each mountain (*i.e.*, skew of the site frequency spectrum) were examined with Tajima’s D^81^, which compares the mean number of pairwise differences against the number of segregating sites observed in a set of sequences, calculated in 50 kb sampling windows for all sites in VCFtools using --TajimaD. We also estimated rarefied private allelic richness per mountain. To this end, we generated per mountain vcf datasets containing only private alleles with BCFtools (view -S -x), which were then merged (BCFtools merge), to create a vcf containing all private variants (1,348,240) in each mountain. This file was converted to a “genind” object where mountains were treated as populations with the R package adegenet^82^. Rarefied private allelic richness was calculated with function allelic.richness in hierfstat, which outputs rarefied allelic counts per locus and population. We used the mean per mountain to visualize private allelic differences between mountains (Supplementary Fig. 1B).

### Demographic and divergence time reconstructions

To better understand temporal changes in demography of East African sky-island *Arabidopsis thaliana*, we used the SMC++ method^50^ to reconstruct past effective population sizes using our dataset of 2,611,238 sites. Reconstructions were carried out independently for each mountain population. We selected a variety of ‘pairs of distinguished lineages’^50^ for each mountain by sampling individuals without replacement and pairing them with individuals sampled with replacement using 12 pairs for Mt. Choke, 14 pairs for Mt. Elgon, 19 pairs for the Simien Mts., and 42 pairs for the Bale Mts. We kept this number close to the number of genetically different individuals per population (based on genetic dissimilarity). We then partitioned vcf files into smc haploblock files for each distinguished pair, further partitioned for each chromosome, and masked the homozygous pericentromeric regions (using genomic coordinates based on TAIR10). Subsequently, we used a polarization error of 0.5 (*i.e.*, the identity of the ancestral allele is not known) and a mutation rate of 7.1 × 10^−9^ (based on results of mutation accumulation experiments^83^), in the ESTIMATE function of SMC++ to estimate past effective population sizes. We ran ESTIMATE by bootstrapping 100 times over 10 randomly selected sequences taken from our dataset of partitioned distinguished pairs. Results were scaled in time using an estimate of 1 yr as generation time and plotted on a linear timescale. To account for seed banking, we also scaled results using 1.5 and 2 yr generation times. Finally, we used these demographies for calculating divergence times between mountains in the cross-coalescent framework of SMC++ in the SPLIT function. The distinguished pairs were determined as described above, but each individual of the pair belonged to a different mountain, using 12 pairs (the minimum number of distinguished pairs in a mountain), resulting in joint frequency spectra for each mountain pair. SPLIT refines the marginal estimates of each mountain into an estimate of the joint demography of mountain pairs. We ran SPLIT by bootstrapping 100 times over 10 randomly selected sequences of the joint SFS.

### Climatic suitability models

We generated a model of climatic suitability in the East African range of *Arabidopsis thaliana* using the maximum entropy framework^84^. We first combined our occurrences (*i.e.*, coordinates at sampled sites) with occurrence data from the Global Biodiversity Information Facility (GBIF, gbif.org) and iNaturalist (iNat, inaturalist.org) both downloaded 20 Dec 2025. We kept GBIF occurrences with coordinates and without flagged problems and iNat observations marked as research grade, resulting in 249 unique locations across the East African range of *Arabidopsis thaliana*. We then used bioclimatic variables that were uncorrelated based on a PCA of climate with 48 CHELSA-BIOCLIM+ variables (Supplementary Fig. 3C) and Pearson *r* < 0.75, including isothermality, temperature of the wettest quarter, precipitation of the driest month, and precipitation of the warmest quarter. These variables are representative of the environmental gradients in the region: isothermality indicates the ratio between warm days-freezing nights over warm summers-cold winters, temperature of the wettest quarter indicates whether rainfall seasons are cold or warm, precipitation of the driest month indicates whether the dry season is severe or mild, and precipitation of the warmest quarter indicates whether there are summer droughts or summer rains. All temperature variables were highly correlated with elevation, with decreasing temperatures at higher elevations across mountains.

For model fitting, we thinned our occurrence points to one sample per 1 km^2^ to reduce sampling bias (N = 105) using the ‘sp’ R package (v.2.2-1)^60^. To characterize potentially inhabited sites, we generated pseudoabsence background points using the ‘dismo’ R package^85^ by sampling 10,000 random points within a 500 km buffer around occurrences^45^. Maximum entropy (MaxEnt) was implemented with the R package ‘ENMeval’ v2, and parameters were optimized using the ‘checkerboard2’ method for cross validation^86^. We chose the model with lowest AICc value and used this to project climatic suitability under recent conditions (1981–2010) across East Africa. For all models, we used the logistic output of MaxEnt that scales suitability from zero to unity. We used permutation importances to determine which climatic factors drove predictions in the distribution model (here temperature of the wettest quarter). We then projected the MaxEnt model using past (LGM) conditions based on the same bioclimatic variables over the past 21,000 years obtained from the CHELSA-TraCE21k-centennial-bioclim dataset^87^.

### Genome-wide associations studies and analysis of allele reuse

We aimed to identified QTL of all phenotypes, including PC1 and PC2 of: germination behavior of seeds from warm-grown plants (Fig. 3A), and growth chamber vegetative and reproductive phenotypes (Fig. 3C). We conducted GWAS that controlled for kinship between genotypes (identity-by-state) using univariate linear mixed models in GEMMA v.0.98.3^88^ on genotypes that had allelic dissimilarity > 0.5 (*i.e.*, with a genome-wide average allele difference greater than 0.5 alleles across pairs of genotypes), excluding loci with MAF < 0.05 for each trait. We corrected Wald test *P*-values using a false discovery rate (FDR) of 0.05 and assigned top SNPs (those with the lowest 100 *P*-values for each trait) to candidate genes using the Araport11 reannotation^64^ and a 50-kb window. We then evaluated elevational allele frequency clines within mountains in GWAS QTL with similar univariate mixed models for GWAS SNP ∼ elevation and a random kinship effect implemented in the R package kinship2 (function lmekin)^89^. Finally, for “warm” phenotypes (overall more variable), we calculated Pearson correlations between mountain pairs’ minor allele frequencies of GWAS SNPs with the 100-top *P*-values. We interpreted *r* > 0.5 as evidence of QTL allele reuse, while negative values indicate significant allelic dissimilarity in QTL.

### Statistical analyses

To understand the potential coordinated multidimensional variation in seed germination behavior, we used PCA in logit transformed germination percentages in each treatment (R function prcomp, variables scaled and centered). To evaluate coordinated multidimensional variation in life history across warm and cold conditions, we used PCA (R function prcomp, variables scaled and centered). To understand how elevational gradients shape phenotypes in each mountain, we tested for mountain-specific phenotype-elevational clines with linear mixed models that controlled for genome-wide similarity among ecotypes. We used the ‘snpgdsIBS’ function in SNPRelate to produce an identity-by-state matrix (from 2.6M genome-wide SNPs in 163 genotypes), used as a random effect. When significant, these models suggest that selection associated with a given phenotype is driving elevational clines, because the cline is stronger than expected by the genome-wide cline^35^. To fit models, we used the function ‘lmekin’ in the kinship2^89^ R package. We focus on mountain-specific elevational clines because across mountains, population genetic structure is hard to control due to strong phenotypic differences between mountain lineages.

## Acknowledgments

Plant material was exported from Uganda with permission of the Ministry of Agriculture, Animal Industry and Fisheries’ Plant Quarantine and Inspection Services, permit UQIS 4414/93/PC (E). Material was exported from Ethiopia with permission of the Ethiopian Biodiversity Institute, Ref. no. EBE71/7065/2018. Material was imported to the USA under USDA APHIS permits P37-17-01651 and P37-18-00230. Funding was provided by NIH award R35GM138300 to JRL. Yuxin Luo, Eleanna Cerda, Haylee Bowers, Margarita Takou, Erica Lawrence-Paul, Shiran Ben-Zeev, Joel Masanga, and Victoria Pizzi assisted in seed bulking and germination experiments. Robin Mailum helped with DNA extractions.

## Data Availability

The sequence data of 148 newly sequenced genotypes generated in this study will be deposited in the European Nucleotide Archive with accession codes available upon publication. The vcf (with 2.6M SNP calls), phenotypes, climate data, and annotated GWAS SNPs will be publicly available via Figshare.

