## Supplementary for "Genomic history and ecology of local adaptation in sky-island *Arabidopsis thaliana*"

This PDF includes

Supplementary Figures 1–8

Supplementary Table 1

### Supplementary Figures

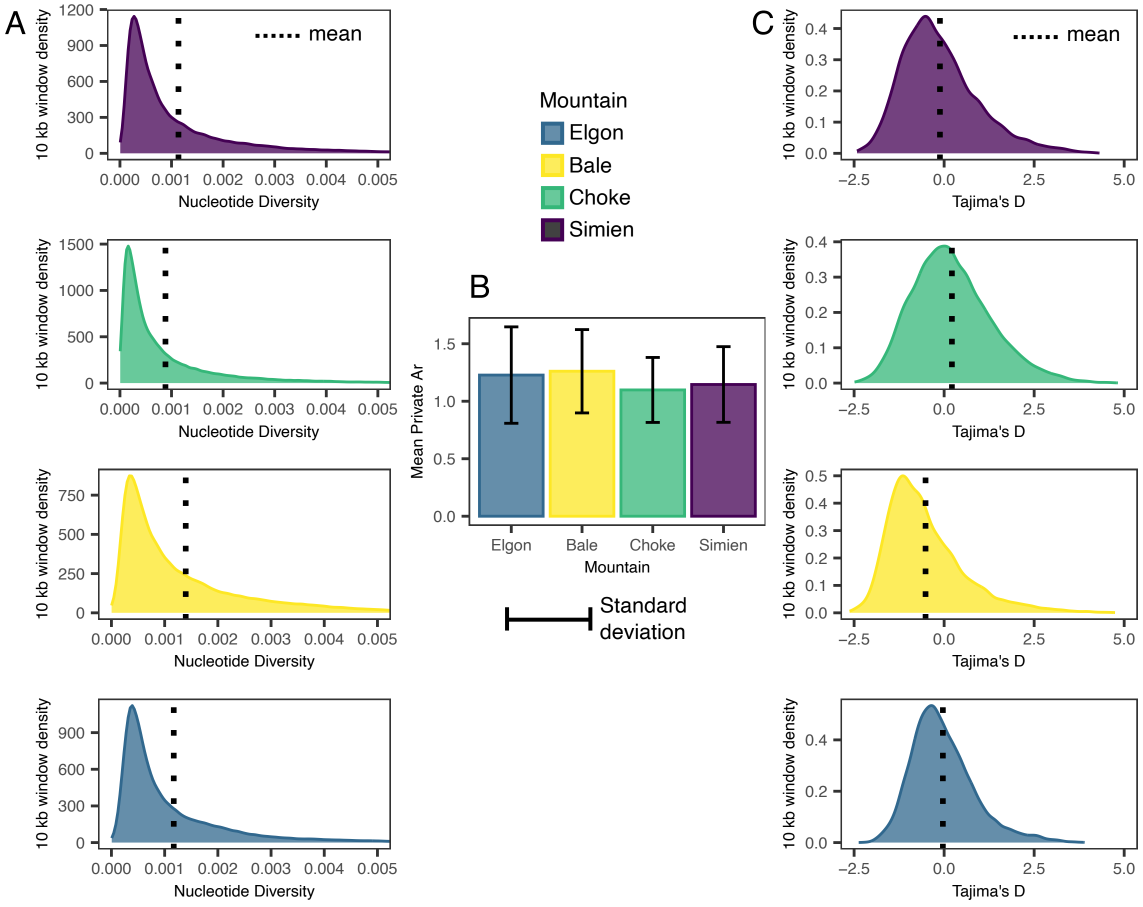


**Supplementary Fig. 1.** Genetic diversity patterns across studied sky-islands. **A.** Genome-wide nucleotide diversity. **B**. Average rarefied private allelic richness. **C**. Skew in the site frequency spectrum (Tajima's D).

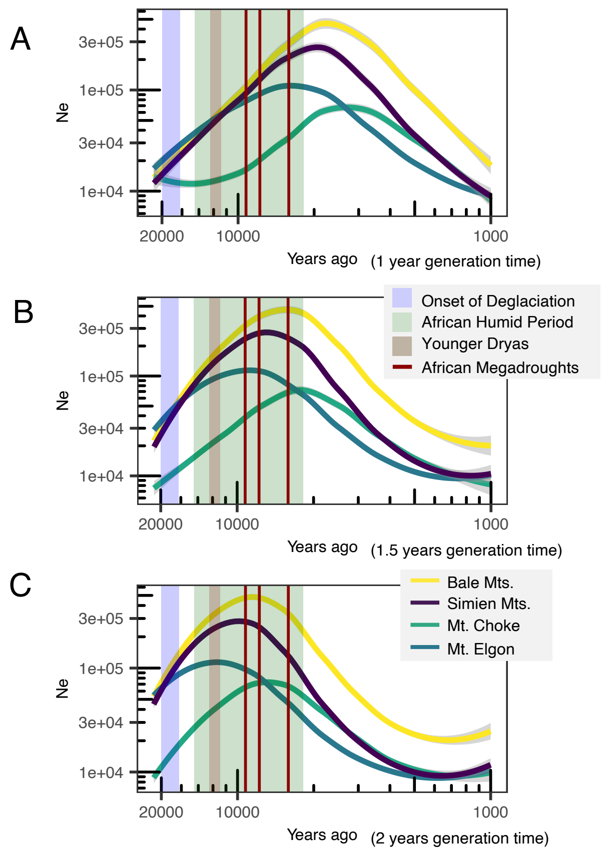


**Supplementary Fig. 2.** *Ne* reconstructions across 20K–1K years ago using different generation times to consider potential seed-banking. **A**. 1-year generation time. **B**. 1.5-year generation time. **C**. 2-year generation time.

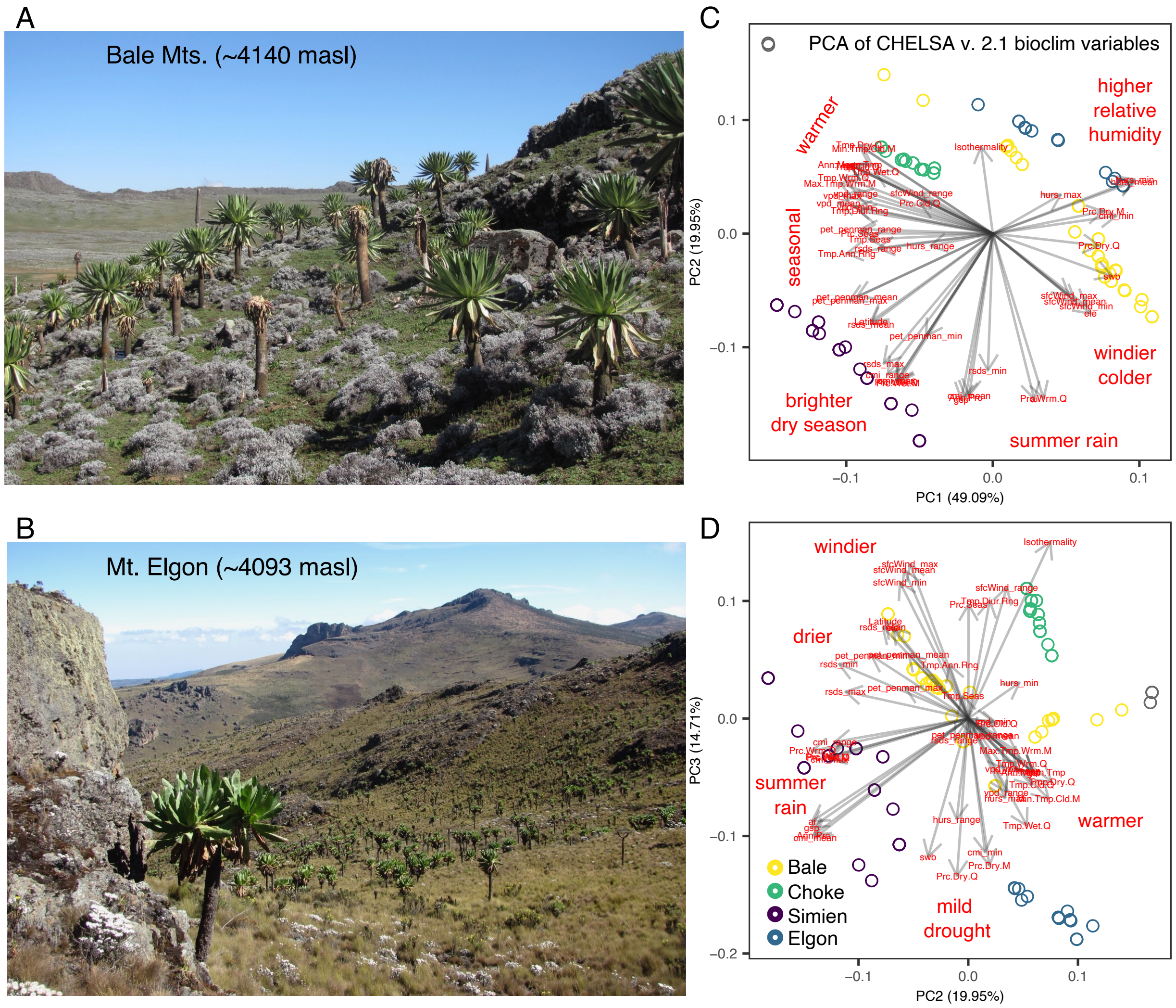


**Supplementary Fig. 3.** Similar habitats and similar elevational gradients, but different climates among studied East African sky-islands. **A**. High elevation site on the Bale Mts. showing the dominant giant rosette *Lobelia rhynchopetalum* (Campanulaceae). **B**. High elevation site in Mt. Elgon, showing the dominant giant rosette *Dendrosenecio elgonensis* (Asteraceae). **C, D**. Principal component 1, 2, and 3 of climate variables from CHELSA v. 2.1 showing studied mountains occupy distinct climates, with the Simien Mts. being the most different.

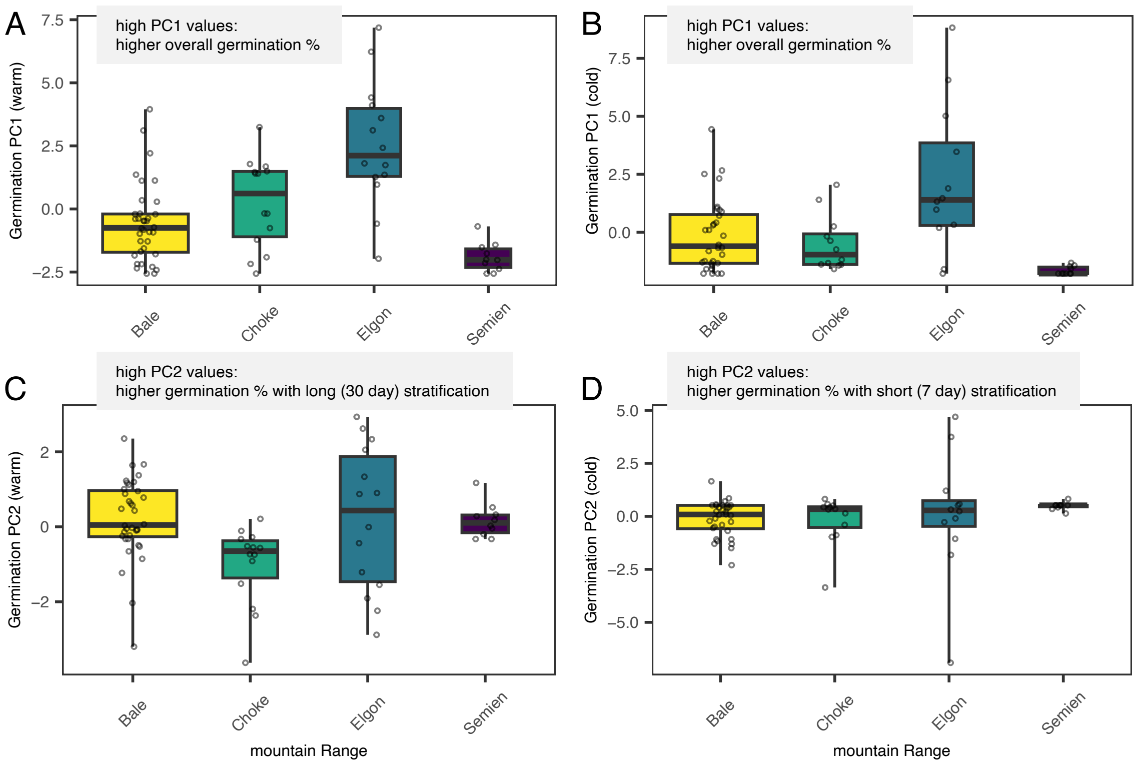


**Supplementary Fig. 4.** Variation within and among mountains in seed germination behavior in seeds from mother plants grown under warm (22ºC day/18ºC night; left) and cool (12ºC day/6ºC night; right) conditions.

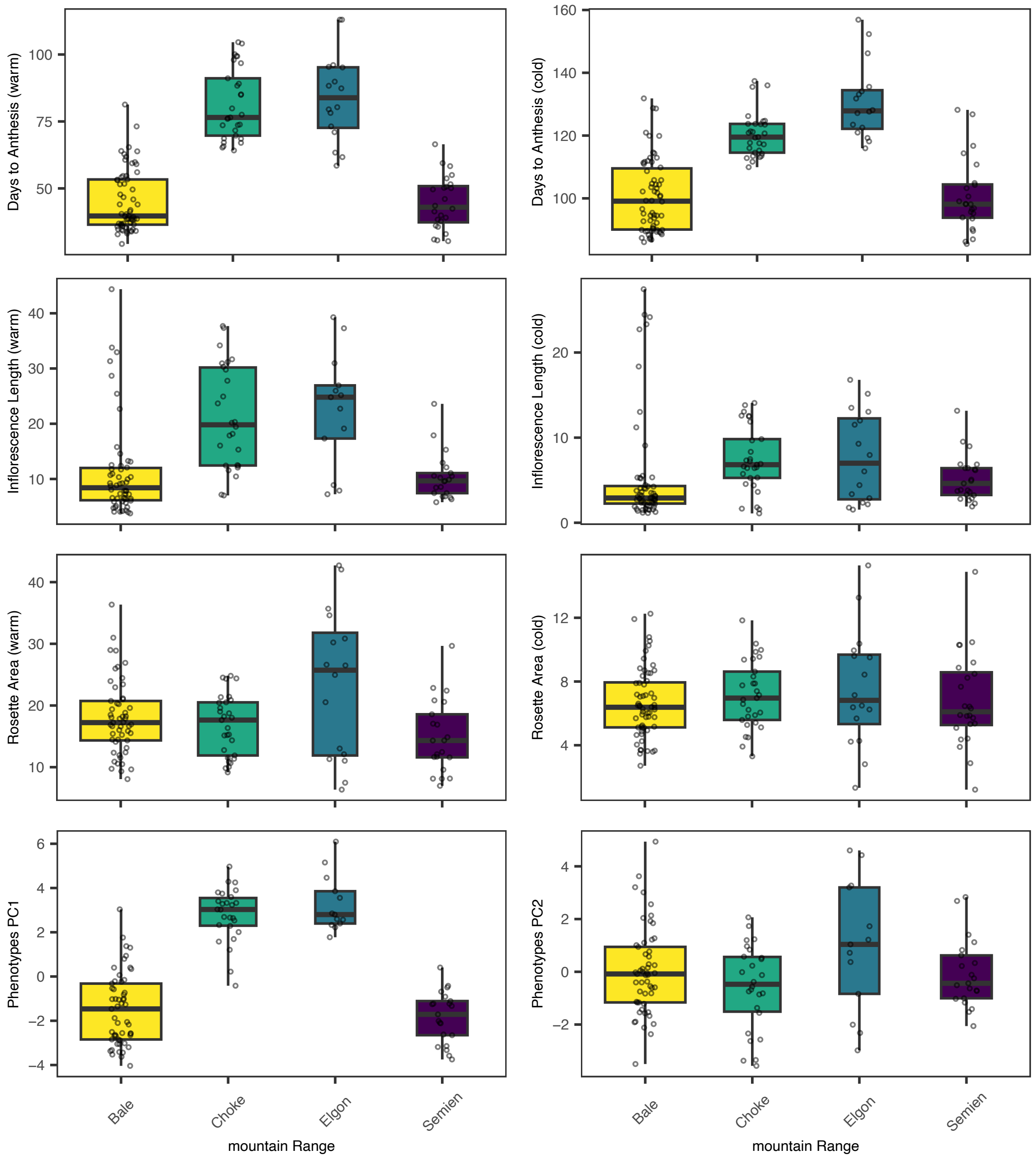


**Supplementary Fig. 5.** Variation within and among mountains in vegetative and reproductive phenotypes measured in warm (22ºC day/18ºC night; left) and cold (8ºC day/4ºC night; right) growth chambers.

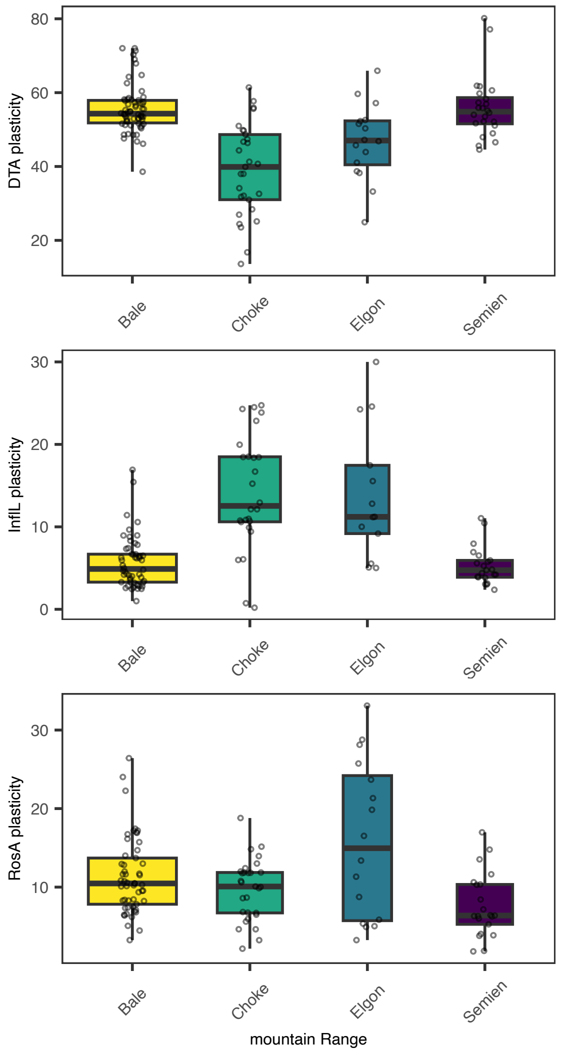


**Supplementary Fig. 6.** Variation within and among mountains in plasticity (trait value under warm – trait value under cold) of vegetative and reproductive phenotypes. Top: Days to Anthesis, middle: Inflorescence Length (cm), bottom: Rosette Area ($cm^2$).

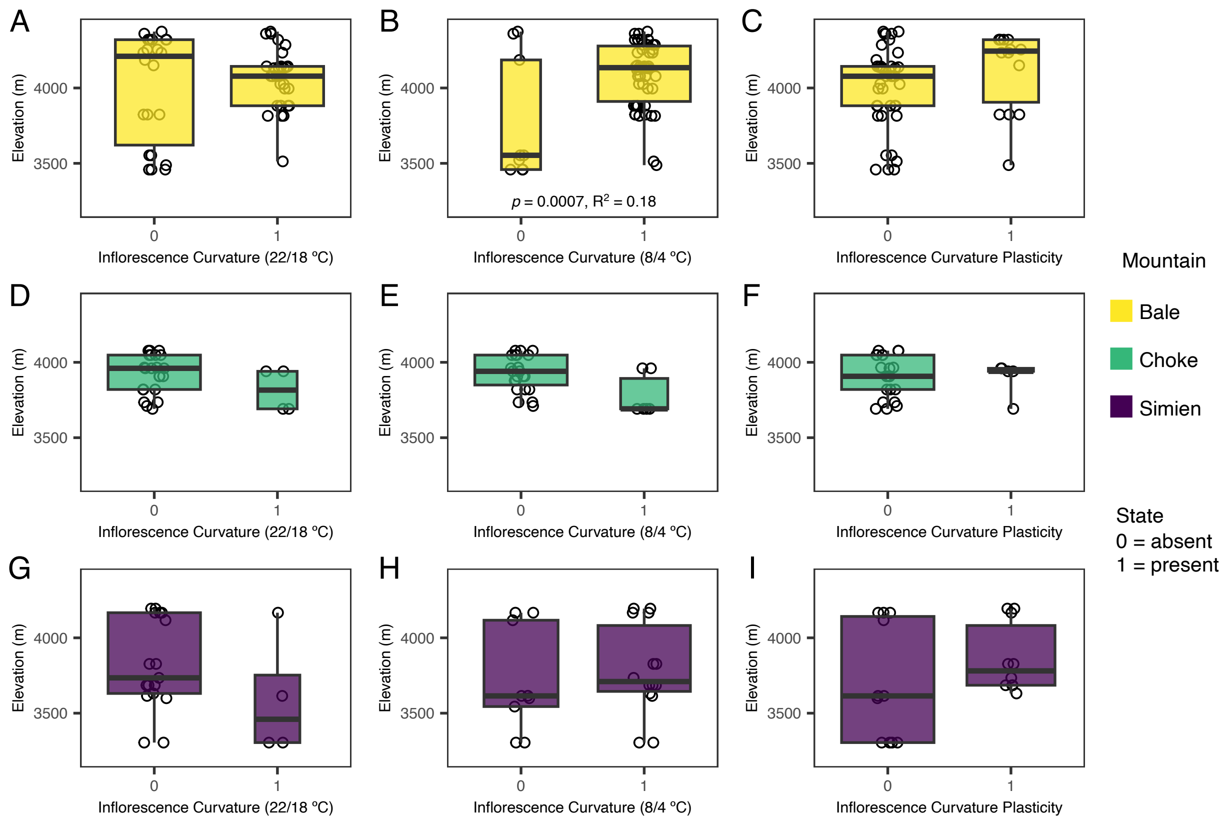


**Supplementary Fig. 7.** Variation along elevation of inflorescence curvature and its plasticity across mountains. Mt. Elgon is not shown here because genotypes there only had erect inflorescences.

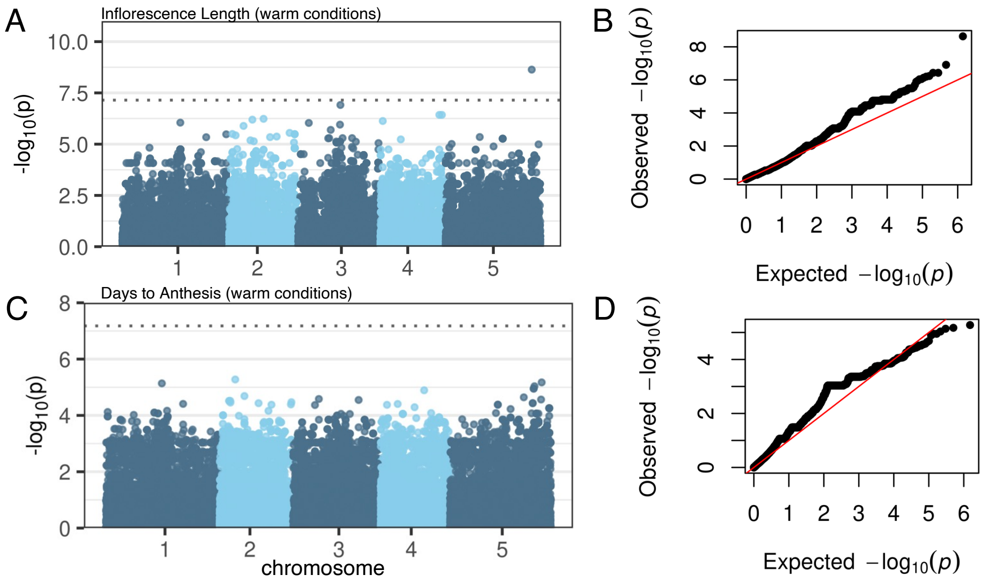


**Supplementary Fig. 8. A, C.** Manhattan plots and **B, D**. QQ-plot of GWAS results for Inflorescence Length (using 66 genotypes and 701,984 SNPs) and Days to Anthesis (using 76 genotypes and 755,492 SNPs), respectively.

**Supplementary Table 1.** Estimates of allele reuse based on Pearson's product-moment correlation between mountain pairs' minor allele frequencies obtained from GWAS SNPs with the 100 top *p*-values for flowering time, inflorescence length, rosette size, and the germination PC1 and PC2 of warm-grown plants.

| Phenotype | Comparison | Pearson *r* | *T* | df | two-side *P* |
|---|---|---|---|---|---|
| Days to Flower | Bale-Choke | -0.471 | -6.51 | 149 | 1.08E-09 |
| Days to Flower | Bale-Elgon | -0.591 | -8.95 | 149 | 1.28E-15 |
| Days to Flower | Bale-Simien | 0.987 | 73.55 | 149 | 2.20E-16 |
| Days to Flower | Choke-Elgon | 0.722 | 12.75 | 149 | 2.20E-16 |
| Days to Flower | Choke-Simien | -0.467 | -6.45 | 149 | 1.45E-09 |
| Days to Flower | Simien-Elgon | -0.575 | -8.57 | 149 | 1.20E-14 |
| Inflor. Length | Bale-Choke | 0.118 | 1.56 | 174 | 1.19E-01 |
| Inflor. Length | Bale-Elgon | 0.674 | 12.05 | 174 | 2.20E-16 |
| Inflor. Length | Bale-Simien | 0.964 | 47.78 | 174 | 2.20E-16 |
| Inflor. Length | Choke-Elgon | -0.122 | -1.63 | 174 | 1.06E-01 |
| Inflor. Length | Choke-Simien | -0.057 | -0.75 | 174 | 4.52E-01 |
| Inflor. Length | Simien-Elgon | 0.744 | 14.69 | 174 | 2.20E-16 |
| Rosette Area | Bale-Choke | 0.988 | 88.18 | 196 | 2.20E-16 |
| Rosette Area | Bale-Elgon | -0.165 | -2.34 | 196 | 0.02 |
| Rosette Area | Bale-Simien | 0.989 | 92.02 | 196 | 2.20E-16 |
| Rosette Area | Choke-Elgon | -0.161 | -2.28 | 196 | 0.02 |
| Rosette Area | Choke-Simien | 0.973 | 59.36 | 196 | 2.20E-16 |
| Rosette Area | Simien-Elgon | -0.143 | -2.02 | 196 | 0.04 |
| Germination PC1 | Bale-Choke | 0.996 | 137.22 | 154 | 2.20E-16 |
| Germination PC1 | Bale-Elgon | -0.938 | -33.54 | 154 | 2.20E-16 |
| Germination PC1 | Bale-Simien | 0.995 | 123.18 | 154 | 2.20E-16 |
| Germination PC1 | Choke-Elgon | -0.936 | -33.10 | 154 | 2.20E-16 |
| Germination PC1 | Choke-Simien | 0.989 | 81.52 | 154 | 2.20E-16 |
| Germination PC1 | Simien-Elgon | -0.944 | -35.45 | 154 | 2.20E-16 |
| Germination PC2 | Bale-Choke | 0.944 | 35.59 | 155 | 2.20E-16 |
| Germination PC2 | Bale-Elgon | 0.606 | 9.48 | 155 | 2.20E-16 |
| Germination PC2 | Bale-Simien | 0.974 | 53.59 | 155 | 2.20E-16 |
| Germination PC2 | Choke-Elgon | 0.610 | 9.58 | 155 | 2.20E-16 |
| Germination PC2 | Choke-Simien | 0.920 | 29.20 | 155 | 2.20E-16 |
| Germination PC2 | Simien-Elgon | 0.593 | 9.16 | 155 | 2.91E-16 |
